# A toolbox for genetic targeting of the claustrum

**DOI:** 10.1101/2024.05.13.593837

**Authors:** Joël Tuberosa, Madlaina Boillat, Julien Dal Col, Leonardo Marconi, Julien Codourey, Loris Mannino, Elena Georgiou, Marc Menoud, Alan Carleton, Ivan Rodriguez

**Affiliations:** Department of Genetics and Evolution, Faculty of Sciences, University of Geneva, quai Ernest-Ansermet 30, 1211 Geneva, Switzerland; Department of Basic Neurosciences, Faculty of Medicine, University of Geneva, 1 rue Michel-Servet, 1211 Geneva, Switzerland

**Author notes:** Corresponding senior authors. Co-first authors (Joël Tuberosa, Madlaina Boillat).

## Abstract

The claustrum (CLA), a subcortical nucleus in mammals, essentially composed of excitatory projection neurons and known for its extensive connections with the neocortex, has recently been associated with a variety of functions ranging from consciousness to impulse control. However, research on the CLA has been challenging due to difficulties in specifically and comprehensively targeting its neuronal populations. In various cases, this limitation has led to inconsistent findings and a lack of reliable data. In the present work, we describe the expression profile of the *Smim32* gene, which is almost exclusively transcribed in excitatory neurons of the CLA and the endopiriform nucleus, as well as in inhibitory neurons of the thalamic reticular nucleus. Leveraging this unique expression pattern, we developed a series of Cre- and Flippase-expressing knockin and BAC transgenic mouse lines with different expression profiles. With these novel tools in hand, we propose new standards for the interrogation of CLA function.

## INTRODUCTION

The CLA is a small, sheet-like, subcortical structure nestled between the insular cortex and the striatum in mammals. Known for its extensive connectivity, it has been described as having the densest reciprocal connections with the neocortex^1–3^, and exhibits anteroposterior and dorsoventral patterning of these connections with different neocortical areas^4–8^. Given its unique connectivity map, the CLA has recently sparked broad interest, and been attributed various functions, including a role in consciousness^9,10^, saliency detection^11,12^, and synchronization of neuronal activity^13^. Recent functional studies, primarily in mice, have suggested roles in a wide range of activities and processes such as behavioral flexibility^14,15^, attention^16,17^, sleep^18–20^, contextual fear conditioning^21–23^, top-down action control^24^, impulse control^25–27^, modulation of cortical activity^28^, pain processing^29–31^, contextual association of reward^32^, and behavioral stress response^33^, among others. While the multifunctional role of the CLA is increasingly recognized, its study has been challenged by difficulties in accurately targeting the structure in murine models due to its complex topology. Unlike the human CLA, which is demarcated by two bundles of white matter fibers, making it clearly distinguishable, the mouse CLA boundaries are unclear: it extends half-way through the brain along the anteroposterior axis with varying spread and cell density^1^, and its demarcation with the insular cortex laterally and corpus callosum medially is not well defined.

To explore CLA function, both historical and contemporary studies have often resorted to direct injections of viruses or tracers into the CLA area. However, these approaches lack cell-type specificity and are, therefore, challenging to replicate.

The use of genetic tools enables precise labeling and manipulation of defined cell populations. This precision is achieved by using specific promoters to drive the expression of transgenes in the desired target cells. To this end, the development of recombinase-expressing mice with specificity in time and space, combined with the use of recombinase-dependent reporter tools, such as viral vectors or transgenic mice, has enabled a wide range of applications. The CLA is primarily composed of projection neurons. The latter express a combination of genes, each showing different degrees of specificity and coverage, such as *Nr4a2*, *Lxn*, *Car3*, *Oprk1*, *Slc17a6* and *Gnb4*^1,15,34,35^. Most of these CLA markers are also expressed in the endopiriform nucleus (EP), ventral to the CLA, and in a subset of cortical layer 6 neurons, rostral and dorsal to the CLA.

A few Cre-driver mouse lines have been used to study the CLA, including Egr2-Cre^17^, Gnb4-IRES2-CreERT2^36^, Slc17a6-IRES-Cre^15^, and Tg(Tbx21-Cre)^18^. Unfortunately, each of these transgenic lines suffers from at least one of the following weaknesses: 1) Cre expression is observed in a poorly defined cell population (in terms of localization or identity) in the CLA area; 2) Cre expression is limited to a subset of CLA projecting neurons (limiting the investigation to this subset); 3) Cre expression requires injection of hormones (potentially limiting behavioral experiments); 4) Cre is expressed in non-CLA populations in the brain in close vicinity to the CLA (specificity is thus difficult to achieve even when using locally-injected viral reporters); and 5) Cre is broadly expressed in the brain and only stops being produced in the area surrounding the CLA during postnatal development (constraining its combination with viral approaches and precluding comprehensive CLA targeting). Overall, establishing a standardized, specific, and exhaustive targeting of CLA cells for its functional interrogation remains challenging today.

Our goal was here to generate genetic tools capable of targeting the majority of mouse CLA projection neurons without affecting other brain cell populations, or tissues outside the brain. Through a search for CLA-specific genes in mice, we identified *Smim32* (a gene with unknown function that is also enriched in the human CLA^37^) as a promising candidate gene to drive recombinase expression in the CLA. Here we present the generation and characterization of different *Smim32* transgenic mouse lines expressing various recombinases in the CLA, providing a new toolbox for researchers aiming at studying the function of the enigmatic structure.

## RESULTS

### *Smim32* is transcribed in the CLA, EN and TRN neurons

We initially searched the entire Allen Mouse Brain Atlas database and visually selected genes with strong and relatively specific signals in the CLA. *Smim32* was found to be highly expressed in the CLA, as well as in the thalamic reticular nucleus (TRN). To precisely determine the identity of neurons expressing *Smim32* in the mouse brain, we re-analyzed published single-cell RNA sequencing (scRNAseq) datasets from dissections of the anterior part of the cortex and the thalamus^38^. Since the full-length mRNA of *Smim32* was not annotated at the time, the analysis involved re-mapping reads using an updated annotation of the mouse genome and reassignment of cellular identities through clustering and manual classification (see Materials and Methods). Among frontal cortex neurons, *Smim32* transcripts were highly enriched in CLA excitatory neurons (Figure 1A-D). These neurons formed a distinct cluster of transcriptomic identities characterized by the expression of *Nr4a2* and *Slc17a6* (Figure 1B,D). Among thalamic neurons, *Smim32* transcripts were highly enriched in inhibitory neurons of the TRN (Figure 1E-H). These neurons also formed a distinct cluster of transcriptome identities characterized by the expression of *Pvalb* (Figure 1F,H).

**Figure 1.**
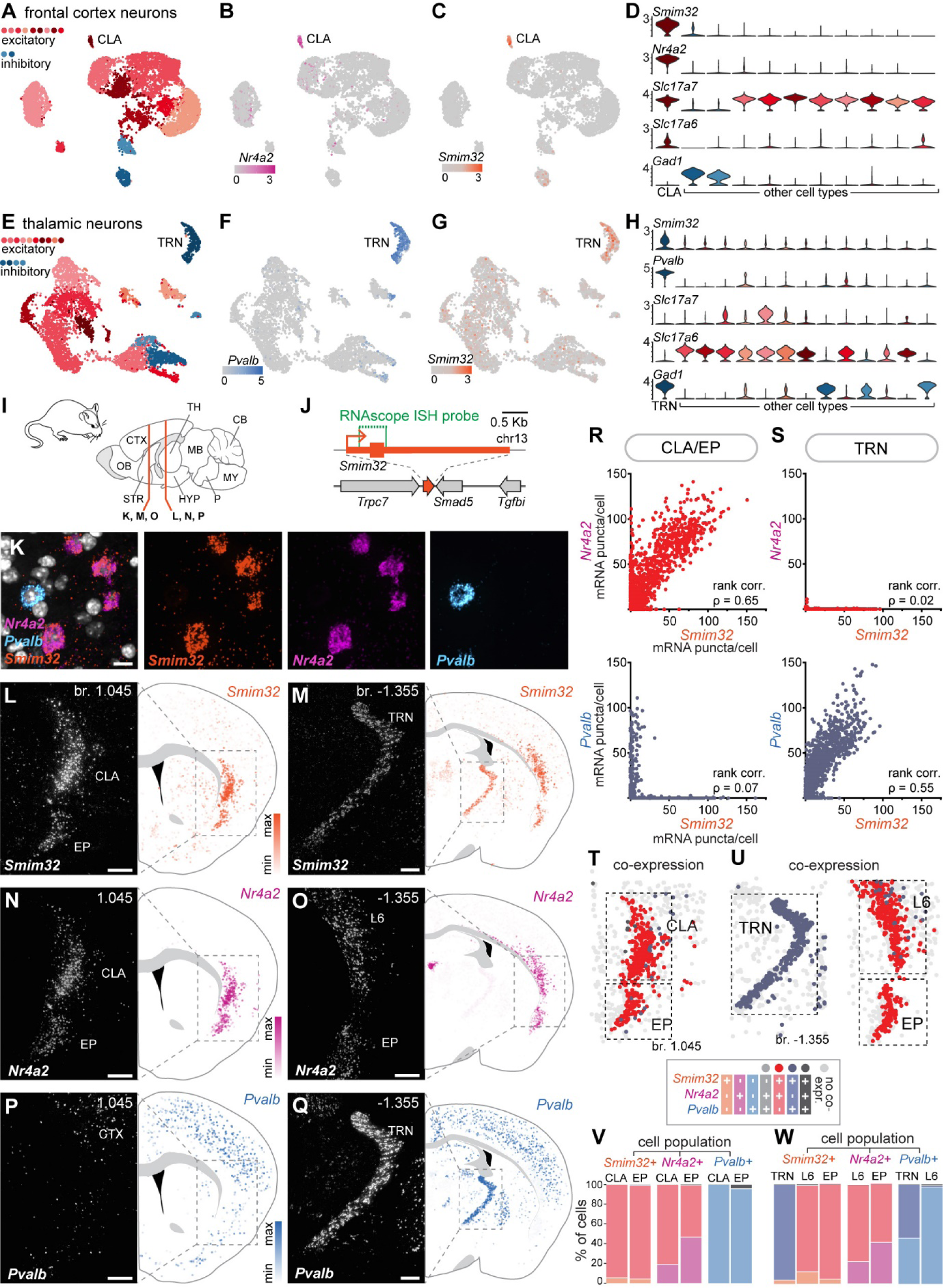
*Smim32* is expressed in the CLA, EP and TRN neurons. (**A**) UMAP plot of single-cell transcriptomes of neurons from the anterior third of the cortex (7804 cells). Excitatory and inhibitory cell types are colored with red and blue tones, respectively. Each cluster is represented by a different color. A single cluster could be identified as excitatory neurons pertaining to the CLA based on the expression of previously identified markers. (**B,C**) Cells shaded according to the normalized expression of (**B**) *Nr4a2* and (**C**) *Smim32* across neurons of the anterior cortex. (**D**) Density plots of normalized expression levels of selected genes across clusters. (**E**) UMAP plot of single-cell transcriptomes of neurons from the thalamus (7317 cells) with a similar color coding as in **A**. (**F**,**G**) Cells shaded according to the normalized expression of (**F**) *Pvalb* and (**G**) *Smim32* across neurons of the anterior cortex. (**H**) Density plots of normalized expression levels of selected genes across clusters. (**I**) Schematic of the localization of coronal sections shown in (**L**-**Q**). Anterior sections (**K**,**M**,**O**) comprise the CLA region, while posterior sections (**L**,**N**,**P**) include the TRN. (**J**) Transcript topology (corresponding to the RefSeq NM_001378296.1) and genomic localization of *Smim32*. The region targeted by RNAscope ISH probes is highlighted in green. (**K**) Illustrative image of CLA cells labeled by RNAscope ISH with probes for *Smim32* (orange), *Nr4a2* (magenta) and *Pvalb* (blue). Individual colored spots (mRNA puncta) correspond to mRNA transcripts. DAPI staining is shown in grey. Scale bar: 10 μm. (**L,M**) Left, representative images of RNAscope ISH labeling of *Smim32* transcripts in the (**l**) CLA/EP and in the (**M**) TRN. Scale bar: 250 μm. Right, cells expressing *Smim32* across the entire hemisection, where dot color corresponds to the *Smim32*-expression level in each individual cell. (**N,O**) Same as above, but showing *Nr4a2* expression in the CLA/EP and in cortical layer 6 (L6). Scale bar: 250 μm. (**P-Q**) Same as above, but showing *Pvalb* expression around the CLA/EP and in in the TRN. Scale bar: 250 μm. (**R,S**) Correlation between expression levels of *Smim32* and *Nr4a2, and Smim32* and *Pvalb,* in cells from the (**R**) CLA/EP and cells from the (**S**) TRN. Each dot corresponds to one cell. All cells from both hemispheres, within the regions of interest (ROI, grey dotted line in (**L,Q**)), were included in the analysis. ρ value indicates Spearman’s rank correlation. CLA/EP, n = 6755 cells; TRN, n = 11139 cells. (**T,U**) Illustrative map of cells co-expressing *Smim32* and *Nr4a2* (red) or *Smim32* and *Pvalb* (purple grey) in the CLA/ EP, the TRN, and in L6. A cell was considered positive for a marker when a minimum of 20 mRNA puncta are attributed to this cell. The dotted lines highlight the ROIs used for quantification analyses shown in (**V,W**). (**V,W**) Bar plots showing the percentages of cells positive for one or several markers. The analysis was done separately for each cell population.

To visualize the expression pattern of *Smim32* in the brain, we labeled *Smim32* transcripts using quantitative RNA in situ hybridization (RNAscope ISH) (Figure 1I-K). Consistent with the scRNAseq analysis, the expression pattern of *Smim32* resembled that of the CLA marker *Nr4a2* within the CLA, EP, and cortical L6 (Figure 1L-O), and of *Pvalb* within the TRN (Figure 1P-Q). To determine the overlap of *Smim32* expression with known CLA and TRN markers, we quantified the expression levels of each marker in individual cells labeled by RNAscope ISH. We confirmed that *Smim32* expression levels correlated with expression levels of *Nr4a2* and *Slc17a6* in the CLA and EP, and with *Pvalb* and *Gad1* in the TRN (Figure 1R-W, Supplementary Figure 1). In the CLA, 94% of all *Smim32* positive cells co-expressed *Nr4a2*, while about 80% of all *Nr4a2* positive cells within the CLA co-expressed *Smim32* (Figure 1V). In the TRN, 96% of all *Smim32*-positive cells co-expressed *Pvalb*, and 54% of all *Pvalb*-positive cells co-expressed *Smim32* (Figure 1W). Thus, *Smim32* is expressed in a large majority of CLA excitatory neurons, as well as in a significant proportion of TRN inhibitory neurons.

### *Smim32* is co-expressed with known CLA markers

We further characterized the expression pattern of *Smim32* within the CLA by examining its potential co-expression with various known markers of the CLA and surrounding insular cortex. RNAscope ISH showed that *Smim32*-expression levels correlated with the expression of *Lxn*, *Gnb4*, *Oprk1,* and *Car3*, all of which are known markers of the CLA (Figure 2A-L). However, *Smim32* was not expressed in cell populations characterized by expression of *Ctgf* (also termed *Ccn2*, cortical layer 6a; Figure 2M-O) or *Rprm* (cortical layer 6b and piriform cortex; Figure 2P-R). Thus, we establish that *Smim32* is a highly specific marker of cells from the CLA and EP in mice.

**Figure 2.**
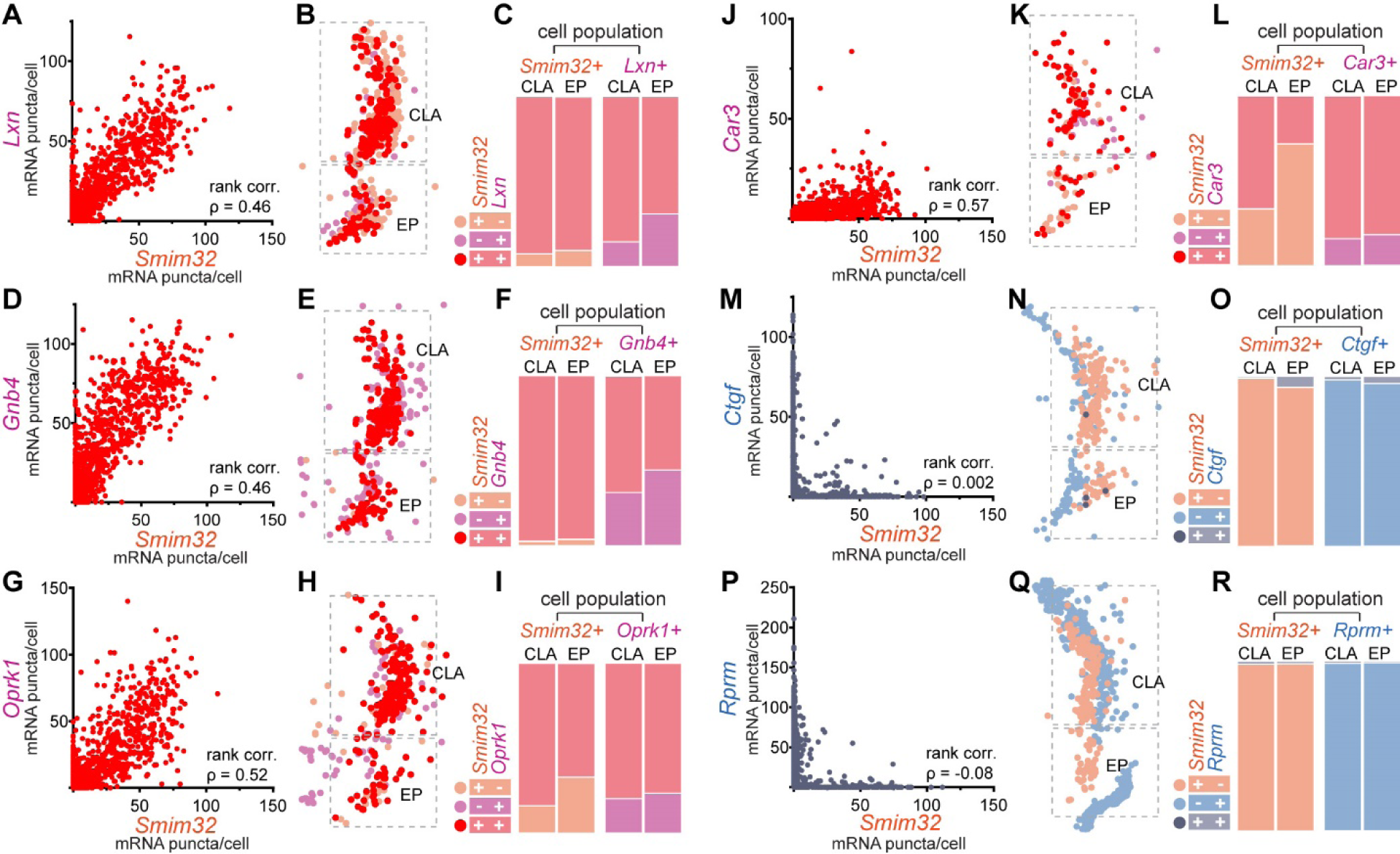
*Smim32* is co-expressed with known CLA markers. (**A**) Correlation between expression levels of *Smim32* and *Lxn* in cells from the CLA and EP. Each dot corresponds to one cell. *Smim32* and *Lxn*: ρ value indicates Spearman’s rank correlation. n = 6429 cells. (**B**) Illustrative plot of cells co-expressing *Smim32* and *Lxn* (red) in the CLA and EP. Dotted lines highlight the ROIs used for quantification analyses shown in (**C**). (**C**) Bar plots showing the percentage of cells positive for one or both markers in the CLA and EP regions. (**D-R**) Same as above, but with different markers of the CLA and surrounding cell populations. (**D-F**) *Smim32* and *Gnb4*, n = 6429 cells. (**G-I**) *Smim32* and *Oprk1*, n = 6780 pairs. (**J-L**) *Smim32* and *Car3*, n = 6493 cells. (**M-O**) *Smim32* and *Ctgf*, n = 7033 cells. (**P-R**) *Smim32* and *Rprm*, n = 6683 cells.

### Generation of mouse lines expressing recombinases in *Smim32*-neurons

To generate mice expressing recombinases in post-mitotic CLA excitatory neurons, we took advantage of the *Smim32* promoter in two separate transgenic approaches. On the one hand, we generated two knockin mouse lines of *Smim32* into which we inserted an IRES-recombinase (either Cre or Flippase) cassette after its endogenous stop codon (Figure 3A). These resulting alleles are later referred to as Smim32-Cre and Smim32-Flpo. To assess the specificity of these tools, we crossed these mice with recombinase-inducible reporter lines. Smim32-Cre mice were crossed with the Cre-inducible tdTomato-expressing reporter line Ai14^39^, and Smim32-Flpo mice were crossed with the Flippase-inducible tdTomato-expressing reporter line Ai65F^39^. Since reporter expression is driven by a CAG promoter and inserted in the Rosa26 locus in both the Ai14 and the Ai65F line, they are both further referred to as R26-tdT. Reporter expression in *Smim32^Cre/wt^*; *R26^tdT/wt^*adult mouse brain was found in a large proportion of the CLA and TRN (Figure 3B-C). Additionally, we observed reporter expression in sparse neurons of the caudoputamen region (Figure 3B) and mid-thalamic region (Figure 3C). The reporter expression pattern was similar in *Smim32^Flpo/wt^*; *R26^tdT/wt^* mouse (Figure 3D-E), except that fewer cells were observed in the mid-thalamic region (Figure 3E).

**Figure 3.**
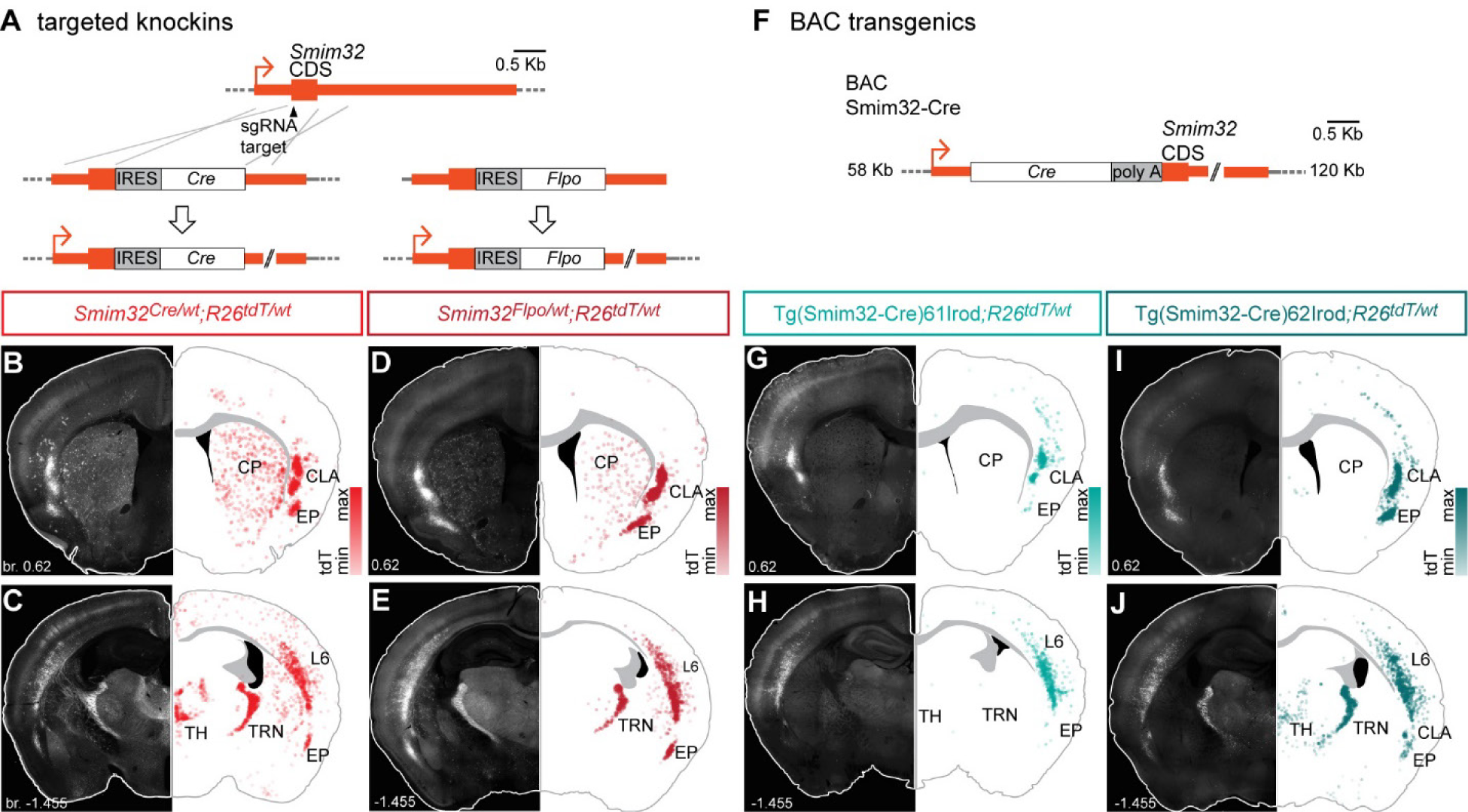
Generation of mouse lines expressing Cre in *Smim32*-neurons. (**A**) Schematic of the CRISPR-mediated insertion of IRES-Cre or IRES-FlpO after the *Smim32* stop codon via homology direct repair (HDR) to generate the *Smim32^Cre^* or the *Smim32^Flpo^* alleles. (**B-E**) Left, epifluorescence images of knockin mice crossed with mice carrying a Cre-dependent or a Flippase-dependent reporter allele *R26^tdT^*. Representative coronal sections at the level of the CLA and TRN. Right, maps of tdT relative expression levels within the represented section. Each dot represents a cell. (**F**) Schematic of the linearized BAC transgene use to generate the different Tg(Smim32-Cre) lines. A detailed description of the transgene can be found in the Materials and Methods section. (**G-J**) Left, epifluorescence images of BAC transgenic mice (**G**,**H**: line 61Irod, **I**,**J**: line 62Irod) crossed with mice carrying a Cre-dependent reporter allele *R26^tdT^*. Representative coronal sections at the level of the CLA and TRN. Right, maps of tdT relative expression within the represented section.

On the other hand, we took advantage of the potential variability in expression between mouse lines associated with classical transgenesis and random transgene insertion in the genome. To this aim, we produced a Cre-expressing transgene under the control of the putative *Smim32* promoter (Tg(Smim32-Cre)) by modifying a BAC containing a genomic fragment spanning the *Smim32* gene (Figure 3F). Successful insertions of this transgene led to different expression patterns observed using the same Cre-inducible reporter as for the Smim32-Cre line (*R26^tdT^*). Two Tg(Smim32-Cre) lines were selected for their specific expression patterns and were further characterized in this work: Tg(Smim32-Cre)61Irod, whose Cre expression is restricted to the CLA (Figure 3G,H), and Tg(Smim32-Cre)62Irod, whose Cre expression is very similar to that of Smim32-Cre, with the exception of a lack of expression in the caudoputamen region (Figure 3I,J).

### Characterization of the expression of reporter genes in adult mice

We then assessed the extent to which transgene-driven reporter expression patterns recapitulated *Smim32*-expression in the knockin and BAC transgenic mouse lines. We quantified single-cell expression of *Smim32* and *tdT* by RNAscope ISH in multiple sections covering different brain areas and at different anteroposterior levels (Figure 4). In *Smim32^Cre/wt^; R26^tdT/wt^* mice, co-expression of *tdT* with *Smim32* reached up to 99.9% in parts of the CLA, indicating highly efficient recombination in these cells and biallelic expression of *Smim32* (Figure 4F). Interestingly, in Tg(Smim32-Cre)61Irod; *R26^tdT/wt^* mice, *tdT* expression was absent in a large fraction of *Smim32*-positive neurons, particularly in the EP. In the TRN of the same line, transgene expression was almost null (Figure 4G), while being expressed in 61-81% *Smim32*-positive cells in the CLA. Tg(Smim32-Cre)62Irod; *R26^tdT/wt^* mice showed a similar reporter expression pattern as *Smim32^Cre/wt^; R26^tdT/wt^* mice (Figure 4H). To evaluate reporter expression in the entire CLA and EP neuronal populations, we examined co-expression of the *tdT* reporter with *Nr4a2* (Supplementary Figure 2). Co-expression between *tdT* and *Nr4a2* reflected the co-expression of endogenous *Smim32* with *Nr4a2* (Figure 1V,W). In the CLA, Tg(Smim32-Cre)61Irod; *R26^tdT/wt^* and Tg(Smim32-Cre)62Irod; *R26^tdT/wt^* mice expressed *tdT* in 60-80% of all *Nr4a2*-positive cells, while in the EP mean co-expression was 23% and 43%, respectively.

**Figure 4.**
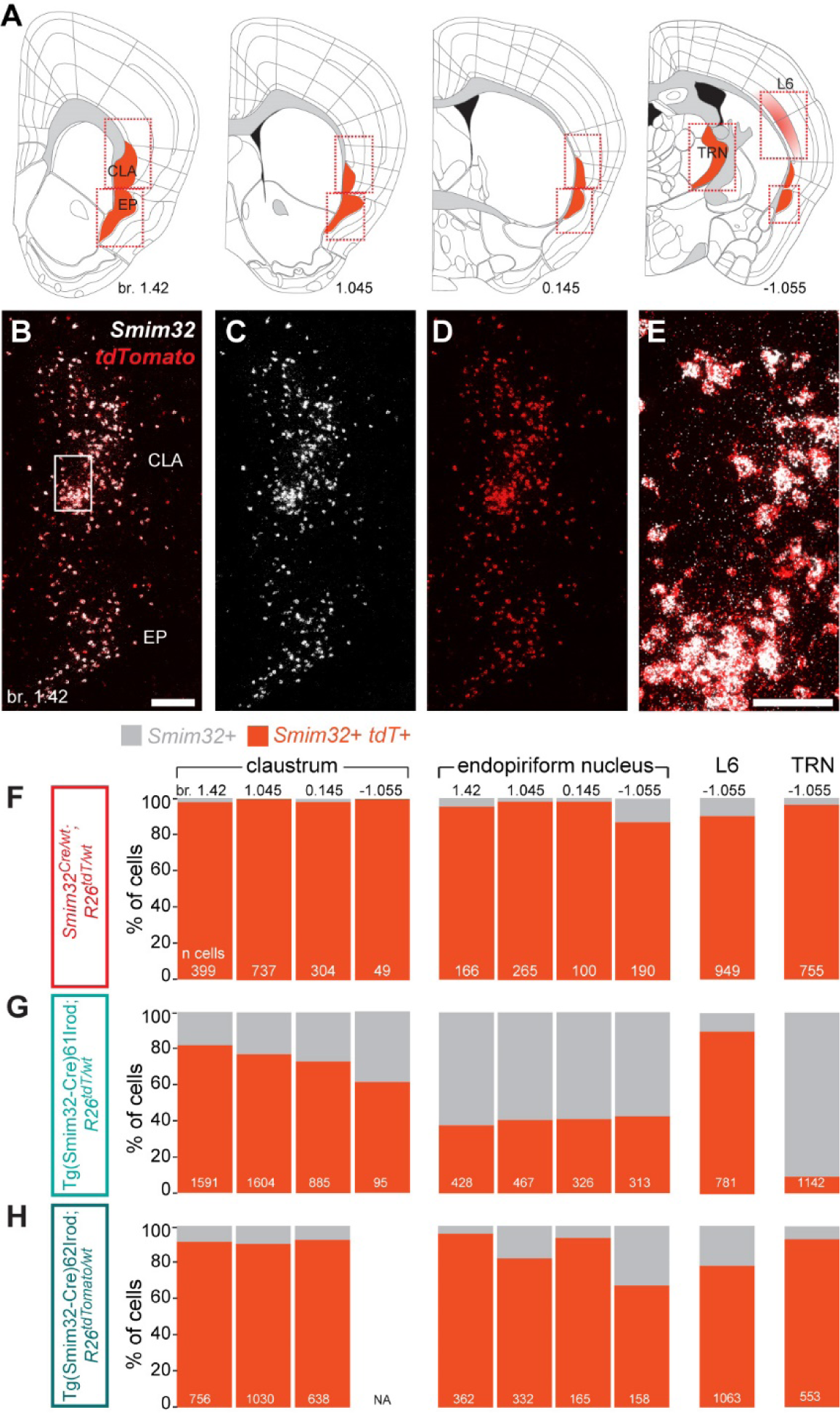
Characterization of the expression of reporter genes in adult mice. (**A**) Schematics showing the regions (according to the Allen Brain Reference Atlases – Adult Mouse, atlas.brain-map.org) that were analyzed to compare expression levels of reporter genes in the knockin and in the BAC transgenic lines. Red dotted lines highlight the ROIs used for quantitative analyses shown in (**F-H**). br: distance from bregma in mm. (**B-D**) RNAscope ISH labeling of *Smim32* (white) and *tdT* (red) mRNA on a coronal section in the anterior CLA of a *Smim32^Cre/wt^;R26^tdT/wt^* mouse. The white square highlights the region magnified in (**E**). Scale bar: 200 μm. (**E**) CLA cells co-expressing *Smim32* and *tdT*. Scale bar: 50 μm. (**F-H**) Bar plots showing the percentage of cells co-expressing *Smim32* with the Cre-inducible reporter allele *R26^tdT^*. A cell was considered as positive for a marker when a minimum of 20 mRNA puncta were attributed to this cell. Numbers in the bar plots indicate the number of cells analyzed in the corresponding region and antero-posterior position.

To further assess the tissue specificity of the two transgenic lines, reporter expression was screened across all organs and tissues of newborn mice of the three Cre-expressing lines. Very sparse reporter-expressing cells were observed outside of the brain, likely resulting from leaky expression of the Cre, with the exception of the kidney medulla and papilla, where reporter-expressing cells were more abundant (Supplementary Figure 3A-C). To evaluate whether these cells expressed *Smim32* in a sustained manner, we re-analysed a published scRNAseq dataset of the mouse collecting ducts^40^. In this dataset, *Smim32* transcripts were detected in no more than 5.2% of each subpopulation (Supplementary Figure 3D), suggesting that the observed patches of reporter-expressing cells result from leaky expression of the Cre and proliferation of recombined cells.

### Characterization of transgenes’ expression during development

To characterize Cre expression patterns of the different Cre lines in space and time, we first evaluated R26-tdT reporter expression in *Smim32^Cre/wt^; R26^tdT/wt^* mice between embryonic day 16 (E16) to postnatal day 200 (P200). The entire brains were sectioned and analyzed, and quantification was carried out in a representative set of coronal levels covering all regions where reporter-expressing cells were found in adult mice (Figure 5A,D, Supplementary Figure 4A,B). For mice aged from E16 to P8, we made a qualitative assessment of *tdT* expression onset (from when the bilateral expression was consistently observed in multiple cells on several consecutive sections). For later ages (P10-P200), we followed the number of reporter-expressing cells across the different regions and time points. Expression started in TRN, thalamus (TH), and main olfactory bulb (MOB), with the first labeled cells visible at E16. In the CLA and EP, expression started at P4. The proportion of cells expressing the reporter was highest in CLA, EP, TRN, and MOB in adult mice, while sparse expression was observed in TH, cortical layers 1-6 (CTX), caudoputamen (CP), and pons. For BAC transgenic mice, we evaluated reporter expression in mice from P0-P70 (Figure 5B-C, Supplementary Figure 4C,D). Expression of transgenes started later compared to the endogenous *Smim32* expression, beginning between P6-P8. Brain-wide analysis of reporter expression confirmed that expression of Cre in Tg(Smim32-Cre)61Irod; *R26^tdT/wt^* mice is highly specific to the CLA, except for sparse expression in the EP and in CTX. In the latter, most of the labeled cells pertained to the layer 6 within posterior sections (see Figure 3G and H for an example).

**Figure 5.**
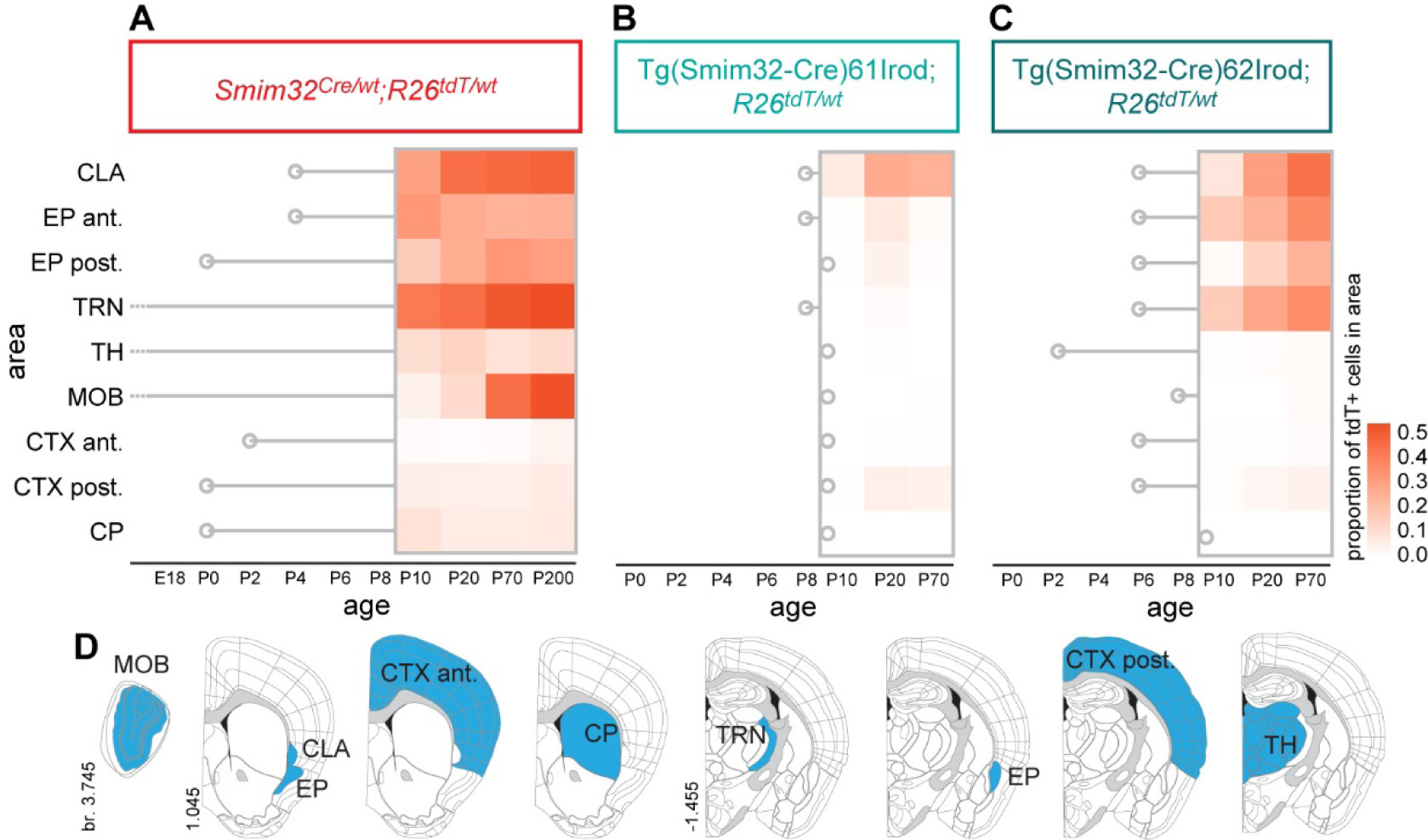
Expression of transgenes during development. (**A-C**) Expression of the Cre-dependent reporter gene *R26-tdT* in brain sections of (**A**) *Smim32-Cre,* (**B**) Tg(Smim32-Cre)61Irod and (**C**) Tg(Smim32-Cre)62Irod mice carrying the *R26^tdT^* allele, from P10 to P200. Only brain regions with persistent expression in adult *Smim32-Cre* mice are shown here. A vertical grey bar indicates the first observation of *tdT*-positive cells within the structure. The heatmap shows relative gene expression levels in sections from P10, normalized across all regions and all mouse lines. (**D**) Schematics of the regions of interest (blue) where *tdT* expression was quantified.

### Cre-inducible tools used with *Smim32* BAC-transgenics

To validate the versatility of the different transgenic Cre-lines generated in this study, we combined our lines with various Cre-inducible genetic tools. Firstly, to test AAV-mediated gene delivery in CLA neurons, we stereotaxically injected a Cre-dependent ChR2-EYFP-expressing AAV2 vector in the CLA region of a Tg(Smim32-Cre)61Irod adult mouse. Three weeks later, immunolabeling of YFP showed restricted recombination in neurons of the CLA (Figure 6A,B). Next, to assess the expression of a different Rosa26 knockin Cre-dependent allele than *R26-tdT*, Tg(Smim32-Cre)61Irod and Tg(Smim32-Cre)62Irod mice were each crossed with the *Gt(ROSA)26Sor^tm^*^2^(CAG–CHRM3*,–mCitrine)*^Ute^/J* (*R26-HA-hM3Dq*) line, which expresses the designer receptor exclusively activated by designer drugs (DREADD) hM3Dq upon Cre recombination (Figure 6C-H). When driven by Tg(Smim32-Cre)61Irod, hM3Dq expression was restricted in CLA excitatory neurons, along with very sparse cells in the caudoputamen and the EP (Figure 6C-D). Furthermore, we evaluated the feasibility of depleting CLA excitatory neurons by crossing Tg(Smim32-Cre)61Irod with the *NSE^DTA/wt^* line, a Cre-inducible diphtheria toxin-expressing reporter transgenic line. Tg(Smim32-Cre)61Irod; *NSE^DTA/wt^* mice displayed an average reduction of 49% of CLA Nr4a2-expressing neurons (Figure I-O). We also examined the expression patterns following BAC transgenic Cre-driven recombination of another family of Cre-inducible alleles, namely Ai155 (TIGRE-Optopatch3) and Ai148D (TIGRE-GCaMP6f). Expression of these reporter genes is controlled by a Tet-responsive element, which relies on a Cre-inducible Tet-transactivator, both of them inserted in the *Igs7* (TIGRE) locus. Tg(Smim32-Cre)61Irod; *TIGRE^Optopatch^*^3^*^/wt^* mice displayed specific expression of the Optopatch3 complex in CLA neurons (Figure 6P,Q), and Tg(Smim32-Cre)62Irod; *TIGRE^GCaMP^*^6f^*^/wt^* displayed specific expression of GCaMP6f in the CLA and TRN neurons (Figure 6R,S). Both reporter expression patterns, respective to the inducing line, are congruent with that of the R26-tdT line under the same condition. In conclusion, the *Smim32*-driven recombinase-expressing lines offer versatile possibilities and reproducibility across different Cre-inducible tools.

**Figure 6.**
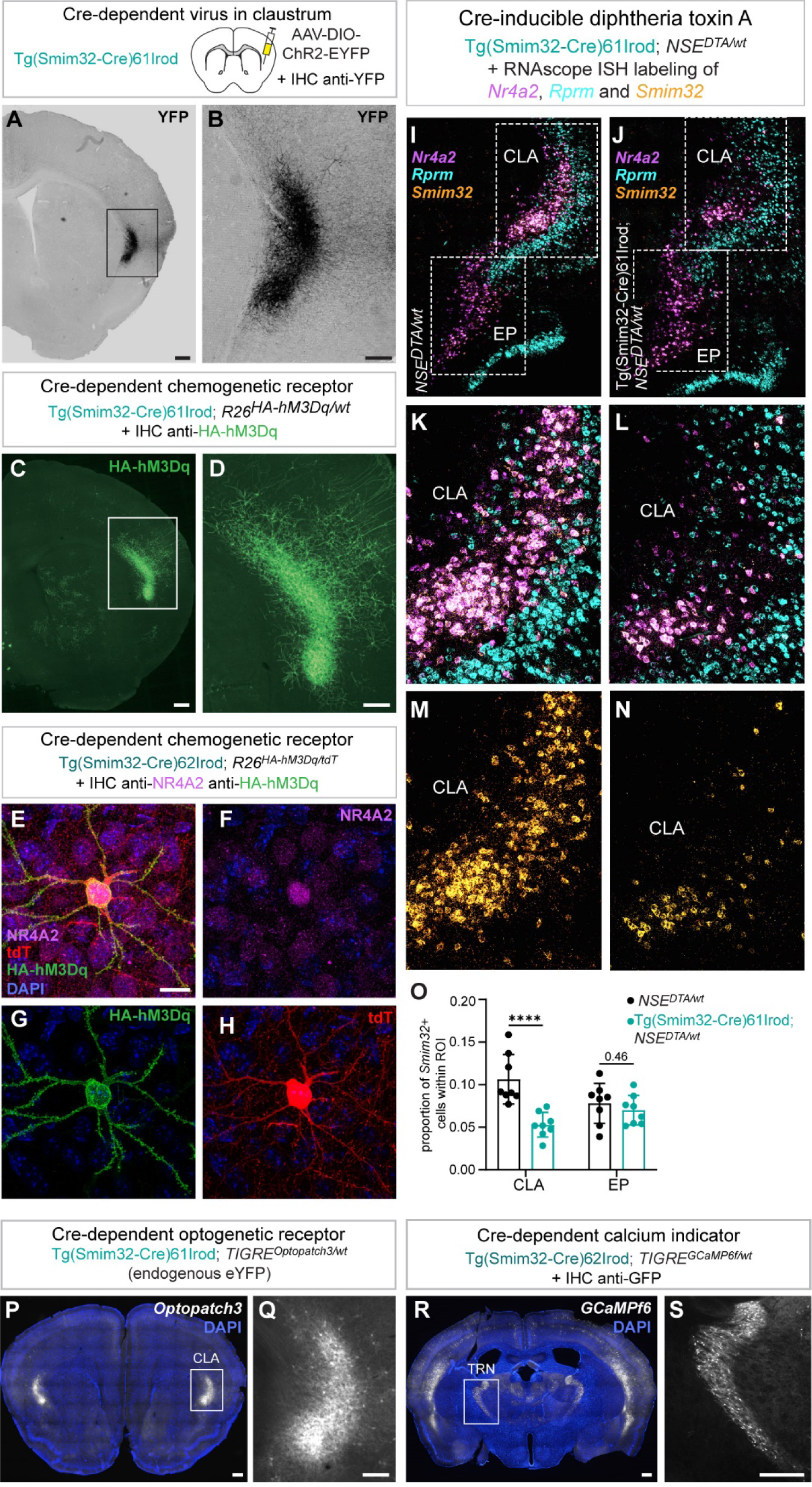
Examples of Cre-dependent reporters used with *Smim32* BAC transgenics. (**A,B**) Tg(Smim32-Cre)61Irod mouse expressing ChR2 and EYFP after injection of a Cre-dependent viral vector with ChR2 and EYFP in the CLA. DAB IHC staining of YFP in a coronal section. The black square indicates the region magnified in (**B**). Scale bars: 200 um (**A**) and 100 um (**B**). (**C,D**) Coronal section of a Tg(Smim32-Cre)61Irod; *R26^HA-hM3Dq/wt^* mouse, where the HA-hM3Dq tag was labeled by IHC (green). The white square highlights the region magnified in (**D**). Scale bars: 200 um (C) and 100 um (D). (**E-H**) Confocal image of a pyramidal neuron in the CLA periphery of a Tg(Smim32-Cre)62Irod; *R26^HA-hM3Dq/tdT^*, DAPI staining of the nuclei (blue), (**F**) NR4A2 immunolabelling (magenta), (**G**) HA immunolabelling (green) and (**H**) tdT endogenous fluorescence (red). Scale bar: 15 um. (**I,J**) Expression of the Cre-inducible diphtheria toxin A in *Smim32*-positive cells of the CLA. Tg(Smim32-Cre)61Irod mice were crossed with *NSE^DTA/wt^* (neuron-specific enolase 2) mice, and the persistence of Smim32-positive cells in the CLA and EP was evaluated by RNAscope ISH labeling. Coronal sections of a Tg(Smim32-Cre)61Irod; *NSE^DTA/wt^* and a control *NSE^DTA/wt^* mouse were labeled with probes against *Smim32* (yellow), *Nr4a2* (magenta) and the cortical layer 6 marker *Rprm* (light blue). (**K-N**) Higher magnification of the CLA region, with all three channels (**K,L**) and only the Smim32 channel (**M,N**). (**O**) Proportion of *Smim32*-positive cells within the CLA and EP (among all cells in the ROI, dotted lines in (**I,J**)). Tg(Smim32-Cre)61Irod; *NSE^DTA/wt^*, n=1 mouse, n= 8 hemisections; *NSE^DTA/wt^*, n=1 mouse, n= 8 hemisections. **** p<0.001, two-way RM ANOVA, Fisher’s LSD test. (**P,Q**) Coronal section of a Tg(Smim32-Cre)61Irod; *TIGRE^Optopatch^*^3^*^/wt^* mouse, with endogenous eYFP fluorescence. DAPI staining of nuclei (blue). The white square highlights the region magnified in (**N**). Scale bars: 200 μm (M) and 100 μm (N). (**R,S**) Coronal section of a Tg(Smim32-Cre)62Irod; *TIGRE^GCaMPf^*^6^*^/wt^* mouse, where GCaMPf6 is revealed by IHC labelling of GFP. DAPI staining of nuclei (blue). Scale bar: 200 μm.

## DISCUSSION

While many groups have attempted to uncover the functions of the CLA, most approaches used to date to target and manipulate the mouse CLA have suffered from an unsatisfactory control or poor description of the cell types studied. In the mouse, neurons with a genetic identity of CLA neurons are nestled in, and even intermingled with other cell populations characterized by the expression of markers such as *Ctgf*, *Nnat*, *Nos1ap* or *Rprm*^15^. Hence, when targeting the CLA by viral injections without the involvement of cell type-specific recombinases, infection of cells from neighboring cell populations is inevitable. This problem cannot be circumvented with dual viral strategies, such as, for instance, the labeling of a subpopulation of CLA projection neurons based on the target area, as the CLA specificity of this approach is not established. Moreover, the use of viruses is often unsatisfactory as the CLA neuronal population is never targeted in its entirety. Unsurprisingly, many approaches essentially based on virus injections have led to apparently contradictory findings and have slowed the emergence of a consensus relative to the various dimensions of the CLA, ranging from its projection pattern to its function.

In this work, we designed a series of recombinase-expressing mouse lines leveraging the promoter of *Smim32*, a gene of unknown function whose expression is specific to CLA and EN projection neurons, as well as to inhibitory neurons of the TRN. These novel Smim32-Cre and Smim32-Flpo alleles provide tools, based on the unique expression pattern of *Smim32*, to specifically target the CLA and EN in the claustro-insular region.

Moreover, taking advantage of peculiar expression patterns resulting from specific integration sites of transgenes, we also generated two additional transgenic lines, Tg(Smim32-Cre)61Irod and Tg(Smim32-Cre)62Irod BAC, which exhibit a more restrictive transcription pattern than the knockin alleles, in particular the lack of expression in the olfactory bulb.

The major advantage of the Tg(Smim32-Cre)61Irod mouse line is its high specificity to CLA neurons: the transgene is expressed on average in 70% of CLA projection neurons (*Nr4a2+* population), while being only sparsely expressed in the EP and other cell populations of the claustro-insular region. Compared to previously described transgenic mouse lines, such as Egr2-Cre^17^, Tg(Tbx21-Cre)^18^, or Gnb4-IRES2-CreERT2^36^mice, which are either expressed in only a small fraction of CLA neurons, or which suffer from high expression in the EP, this mouse allows the specific targeting of a large fraction of CLA neurons.

Taking advantage of the new Smim32-FlpO allele, even more specific targeting of CLA projection neurons could be achieved via an intersectional approach, involving, for instance, Slc17a6-Cre and Smim32-FlpO. Using the desired reporter, this will allow broad characterization of CLA projections, specific FACsorting of CLA neurons, optogenetic and pharmacogenetic manipulation of the whole structure, without any virus delivery.

With these genetic tools in hand, the comprehensive targeting and functional probing of the densely connected CLA is now possible.

## MATERIALS AND METHODS

### Animals

All experiments were conducted in accordance with the veterinary guidelines and regulations of the University and of the state of Geneva (Direction of Animal Experimentation, UNIGE). C57BL/6J male mice were purchased at 5–7 weeks of age from Charles River Laboratories. Upon arrival, they were group-housed in standard type II cages with *ad libitum* access to food and water. They were held on a dark/light cycle of 12/12 h, temperature between 21 and 22 °C, and humidity between 45 and 55%.

### Generation of transgenic mice

#### BAC transgenics Tg(Smim32-cre)61Irod and Tg(Smim32-cre)62Irod

This transgene was obtained by a modification of the bacterial artificial chromosome (BAC) RP23-125G21, from the RPCI-23 BAC library^41^. This BAC spans from the 3’ intergenic region in 5’ of *Smad5* to the second intron of *Trpc7*, therefore encompassing *Smim32*, which is located in between these two genes. To generate this transgene, we inserted the cassette DDCre-Rabbit β-globin polyA-frt-PuroR-ZeoR-frt in place of *Smim32* start codon via recombineering. The resulting vector was injected in the pronuclei of C57BL6/J-DBA/2J F2 zygotes. Four founders were obtained out of which two lines were established for their expression pattern specificities, hereafter named Tg(Smim32-cre)61Irod and Tg(Smim32-cre)62Irod. DDCre refers to the fusion of a destabilizing tag at the N-terminus of the Cre recombinase, targeting the protein to proteasome degradation^42^, a process that is inhibited by trimethoprim (TMP) treatment. Three intraperitoneal injections of 300 µg of TMP/gram of body weight on three consecutive days one week prior to analysis (one-month-old animals) did not affect the reporter (R26-tdT) expression pattern in either transgenic line. For this reason, we removed the DD prefix from our nomenclature.

#### Smim32-Cre

This knockin was obtained via CRISPR-induced homology direct DNA repair (HDR) using a circular donor template. The vector of this template is a pUC57 plasmid in which we inserted an IRES-Cre (Cre) cassette flanked by homology arms corresponding to 801 bp and 799 bp before and after the cut site, respectively. The expected cut site, which is also the insertion site, was located on the *Smim32* stop codon. The HDR donor along with the Cas9 complex (Cas9NLS protein, tracrRNA and crRNA) was injected in the pronuclei of C57BL6/J mice. A unique founder was selected and backcross for at least three generations before producing the mice analyzed in this study.

The guide RNA target sequence is AGACGCGACCGGTATAGGGC(AGG) (PAM sequence in parentheses).

#### Smim32-Flpo

This knockin was obtained via CRISPR-induced HDR using a Tild vector^43^. The guide RNA target is the same as for Smim32-Cre. The targeting vector was modified from used for Smim32-Cre, with slightly shorter homology arms resulting from blunt digestion (796 bp and 797 bp in 5’ and 3’, respectively) and the codon-optimized Flippase (FlpO) coding sequence in place of the Cre. The donor template (a linear fragment composed of the aforementioned sequences) was injected along with the Cas9 complex in the pronuclei of C57BL6/J mice. A unique founder was selected and backcross for at least two generations before producing the mouse analyzed in this study.

### Genotyping

Tg(Smim32-cre) mice can be genotyped by PCR with the primers #1 and #2 (see Table 1), yielding a 854 bp amplicon from the transgenic allele. *Smim32-Cre* mice can be genotyped with a mix of the primers #3, #4 and #5, yielding a 400 bp amplicon from the *Smim32-Cre* allele and a 645 bp amplicon from the wild-type allele. *Smim32-Flpo* mice can be genotyped with a mix of the primers #3, #5 and #6, yielding a 423 bp amplicon from the *Smim32-Flpo* allele and a 645 bp amplicon from the wild-type allele.

**Table 1.**
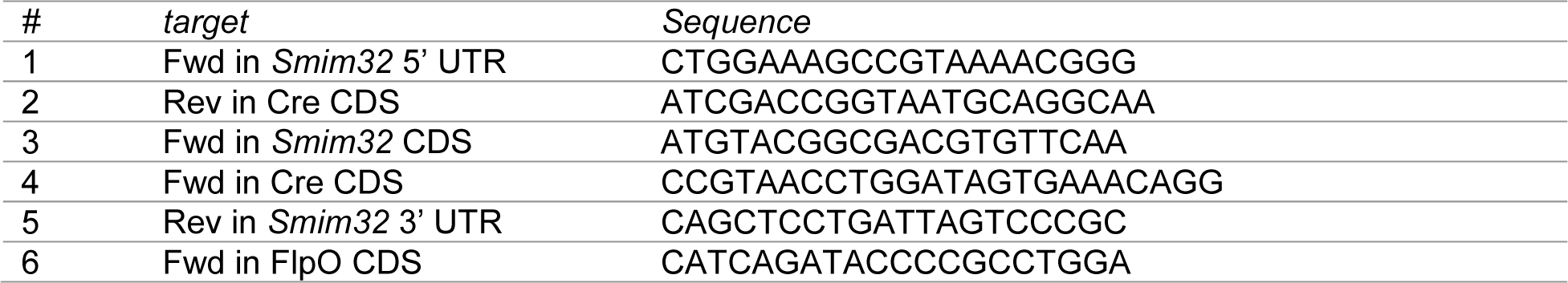
Primers for genotyping.

### Reporter Mouse lines

Details regarding all transgenic mouse lines used are provided in Table 2.

**Table 2.**
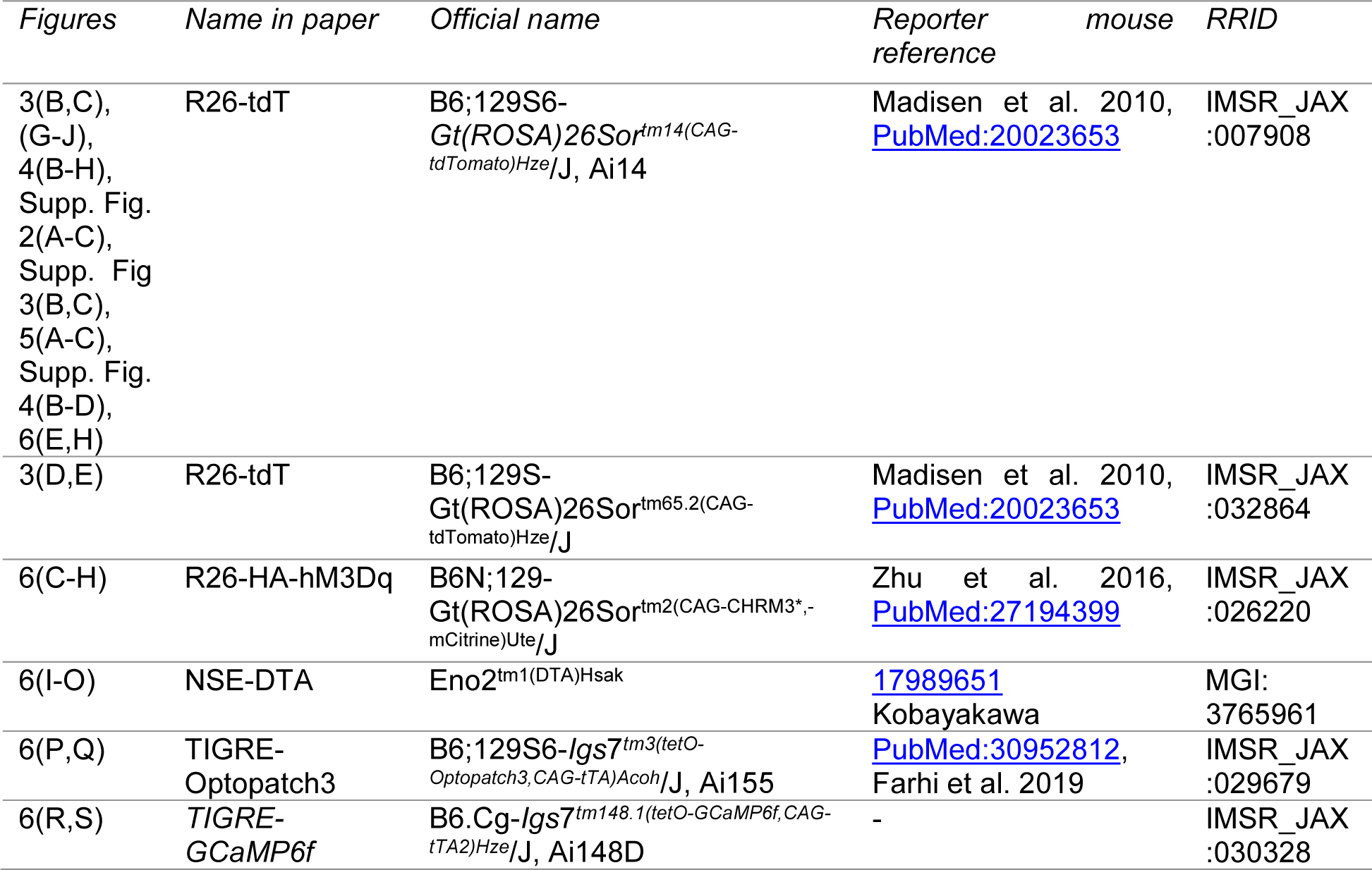
List of transgenic mouse lines.

### Single-cell RNAseq analysis

#### Brain

Single-cell RNA-sequencing (scRNA-seq) datasets from the Linnarson lab’s mouse brain atlas (http://www.mousebrain.org^38^) corresponding to the anterior part of the cortex and to the thalamus were downloaded from NCBI’s sequence read archive (SRA, see run accessions in Table 3).

**Table 3.** SRA run accessions for single-cells RNAseq analysis.

| *sample identifier* | *tissue* | *SRA run accession* |
| --- | --- | --- |
| 10X05_1 | anterior cortex | SRR6854100 |
| 10X05_2 | anterior cortex | SRR6854145 |
| 10X22_1 | anterior cortex | SRR6854092 |
| 10X22_2 | anterior cortex | SRR6854170 |
| 10X24_3 | thalamus | SRR6854165 |
| 10X24_4 | thalamus | SRR6854164 |
| 10X50_5 | thalamus | SRR6854143 |
| 10X50_6 | thalamus | SRR6854144 |
| 10X50_7 | thalamus | SRR6854137 |
| 10X50_8 | thalamus | SRR6854138 |
| 10X57_6 | thalamus | SRR6854074 |
| 10X57_7 | thalamus | SRR6854064 |

*Smim32* 3’UTR was partially annotated at the time of the publication of Linnarson lab’s mouse brain atlas^38^and, therefore not detected. To capture *Smim32* expression levels in these data, we updated the Ensembl annotation version 86 of the mouse genome assembly mm10/GRCm38 with the annotation of the RefSeq mRNA NM_001378296.1, which exhaustively covers the 3’ UTR. Extraction of the raw data and mappings on the updated mouse genome annotation were performed with the cellranger software suite, provided by the 10X Genomics website (https://support.10xgenomics.com/). Sequences were extracted from the BAM files using the program bamtofastq, and mapping and counting were performed with the count program, that was run using the options --force-cells=3500. The clustering analysis was performed using the Seurat package^44^. First, single cell transcriptomes (thalamus: 37000 cells from height samples, anterior cortex: 14000 cells from four samples) were filtered to remove datasets with less than 600 counts or with an insufficient ratio of UMI per number of genes (nGene) detected (UMI/nGene >= 1.2). Next, genes that were present in less than 20 cells were removed from the analysis. The samples were merged and integrated using the SCTransfom protocol, available on the Seurat website (https://satijalab.org/seurat/v3.1/integration.html). Clusters were formed using the findneighbours and findcluster functions from Seurat with the principal components (anterior cortex: PC 1-8, resolution 0.2, thalamus: PC 1-14, resolution 0.2). Next, we reduced the dataset to neurons by selecting cells expressing the neuronal marker *Snap25*. A second integration was performed on this subset using the same protocol as for the whole dataset, but with slightly different parameters (anterior cortex: PC 1-8, resolution 0.1, thalamus: PC 1-13, resolution 0.2). Finally, a dimensionality reduction was performed with the UMAP method to project the transcriptomes on a 2-dimensional plot (Figure1A-D).

#### Kidney

Mouse kidney collecting duct scRNAseq data was retrieved from NCBI’s SRA under the accession SRP108643. Mapping on the mouse genome assembly mm10 was performed with STAR^45^, restricting to uniquely mapped reads with the option --outFilterMultimapNmax=1. Quantification of *Smim32* expression was performed with featureCounts^46^ using the same updated gene annotation file as for the brain scRNAseq data. Cell type annotation was retrieved from the corresponding GEO dataset GSE99701. To estimate the fraction of *Smim32*-expressing cells across cell types (pie charts in Supplementary Figure 3D), we counted the number cells with *Smim32* transcript levels reaching more than 1 TPM over the total number of cells.

### RNAscope *in situ* hybridization

Mice were anesthetized with isoflurane and killed by cervical dislocation, brains were immediately embedded in OCT and frozen in an isopentane bath in liquid nitrogen. Samples were stored at - 80°C until sectioning. Brains were cut on a cryostat microtome in 10 μm coronal sections and mounted on Superfrost Plus slides. Sections were dried for 1-2 hours at -20°C inside the cryostat, before being transferred for storage at -80°C. Individual mRNA molecules on sections were labeled using the RNAscope™ Multiplex Fluorescent V2 Assay (ref. 323136, Advanced Cell Diagnostics), following the guidelines for fresh-frozen tissue. Before probe hybridization, sections were post-fixed in Formalin 10% for 15 min at 4°C, dehydrated, treated with hydrogen peroxide for 10 min at room temperature, and treated with Protease IV for 15 min at room temperature. Probes were visualized with TSA Vivid Fluorophores 520 (1:750), ref. 323271, 570 (1:1500) ref. 323272 and 650 (1:2000), ref. 323273, diluted in TSA buffer. Sections were counterstained with DAPI and mounted with ProLongTM Gold antifade (ref. P36935, Invitrogen). Slides were imaged with a Nikon Ti/CSU-W1 Spinning Disc Confocal microscope using 405, 488, 561 and 640 nm excitation lasers and a 60x 1.49 NA air objective. Image tiles of 6 Z planes cover 4 μm were acquired, merging of tiles and orthogonal projection was applied within the SlideBook software (Intelligent Imaging Innovations).

### RNAscope probes

All RNAscope probes used in the present study can be found in Table 4.

**Table 4.**
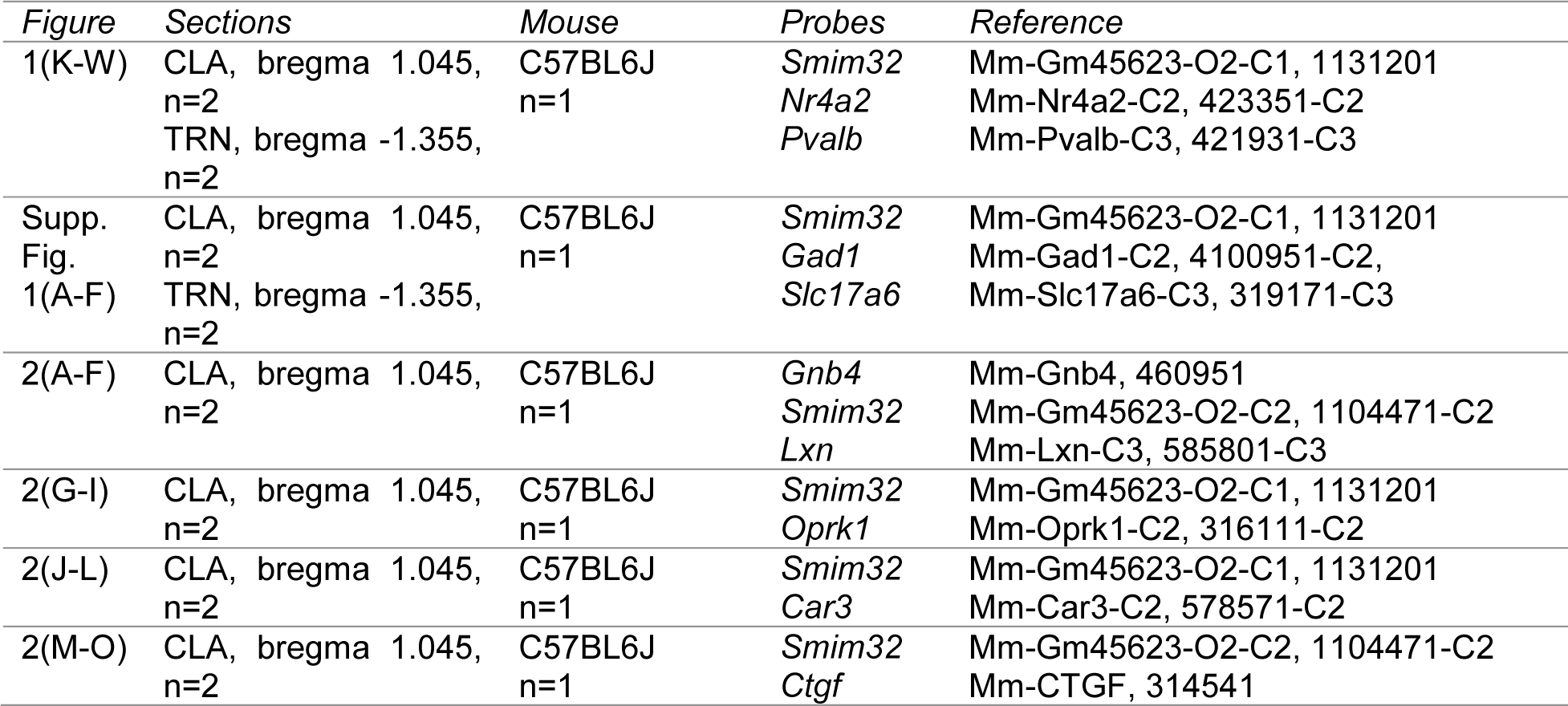
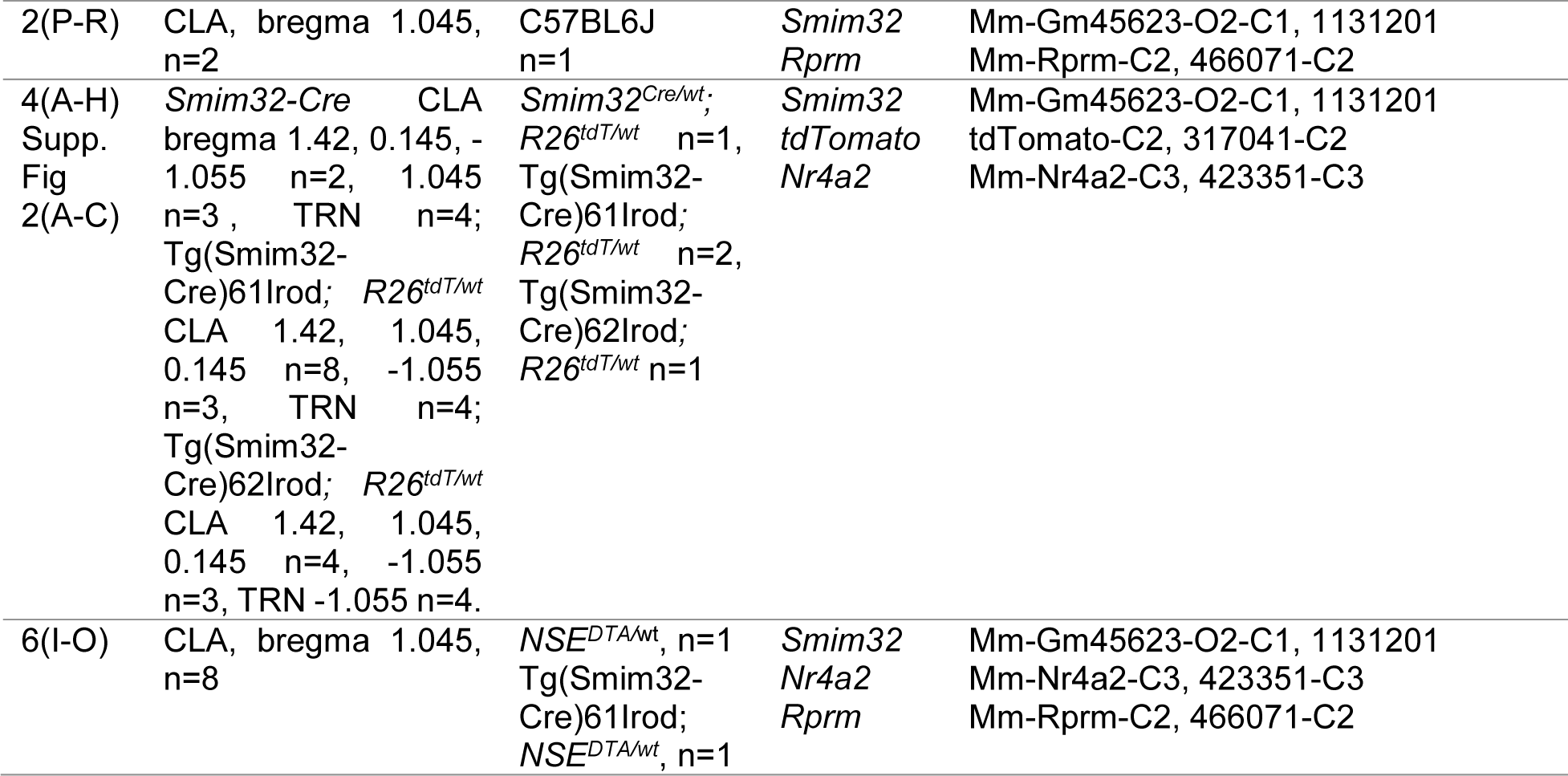
List of RNAscope probes.

**Table 5.**
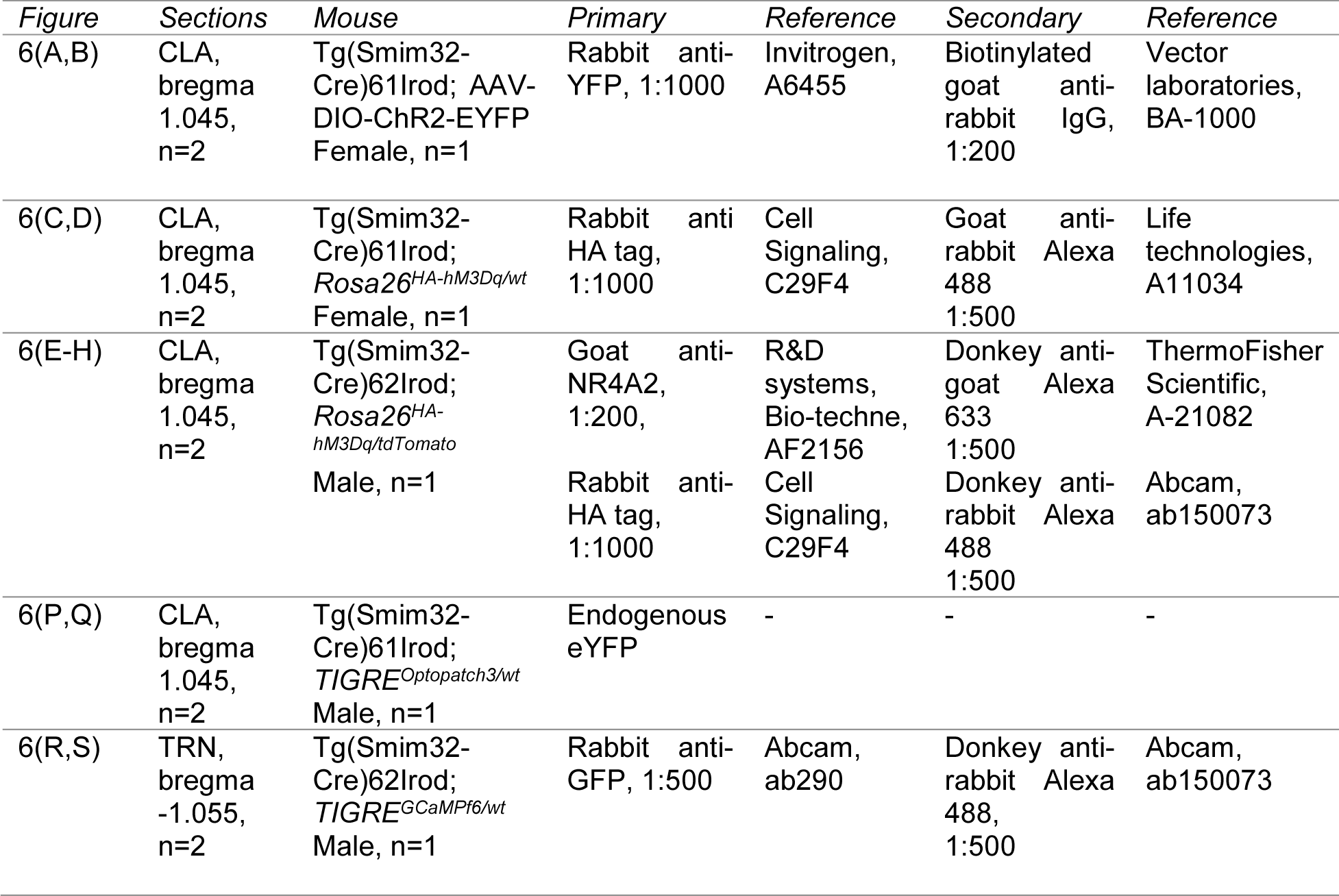
List of antibodies used for IHC labeling.

**Table 6.**
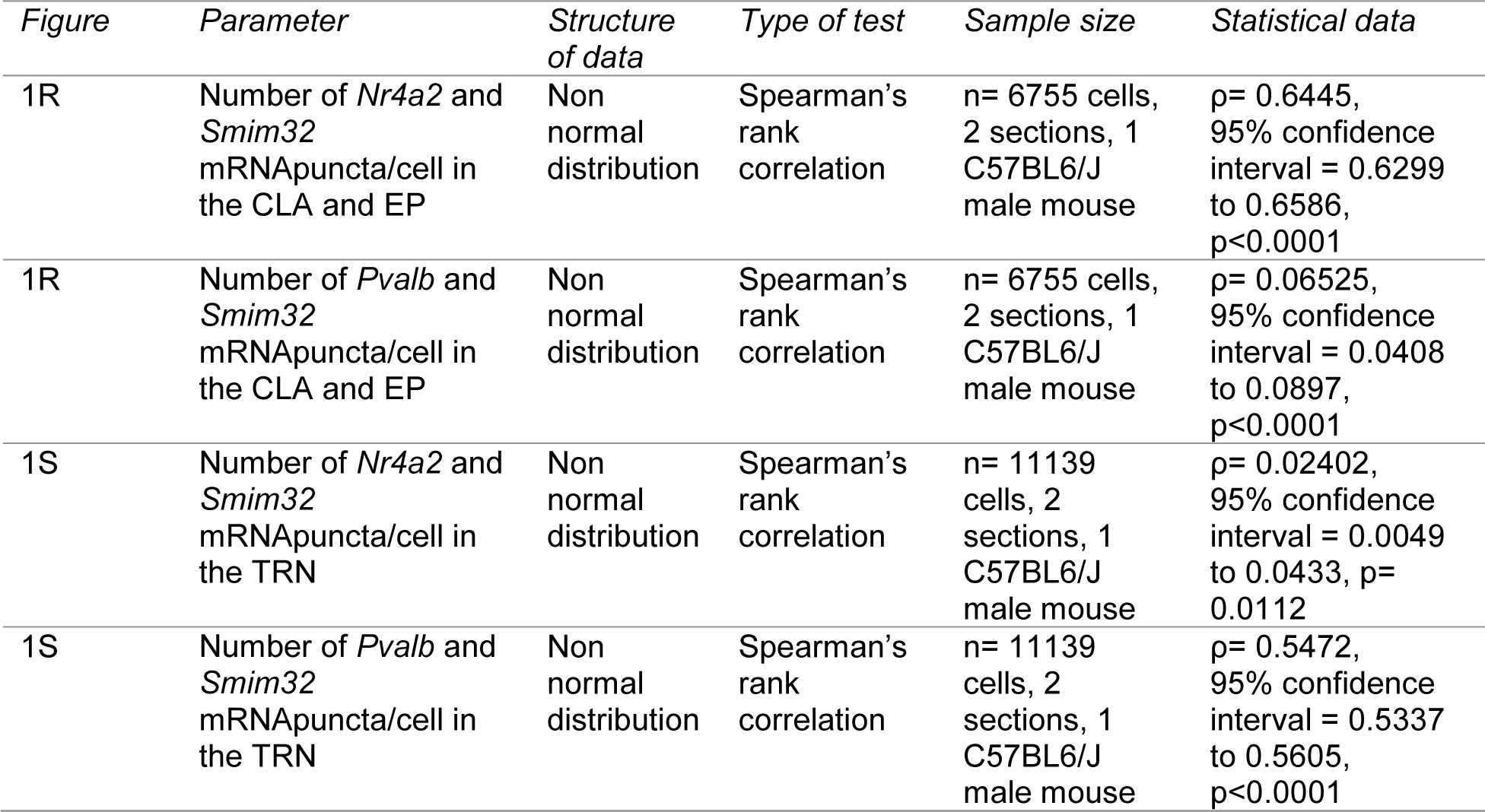
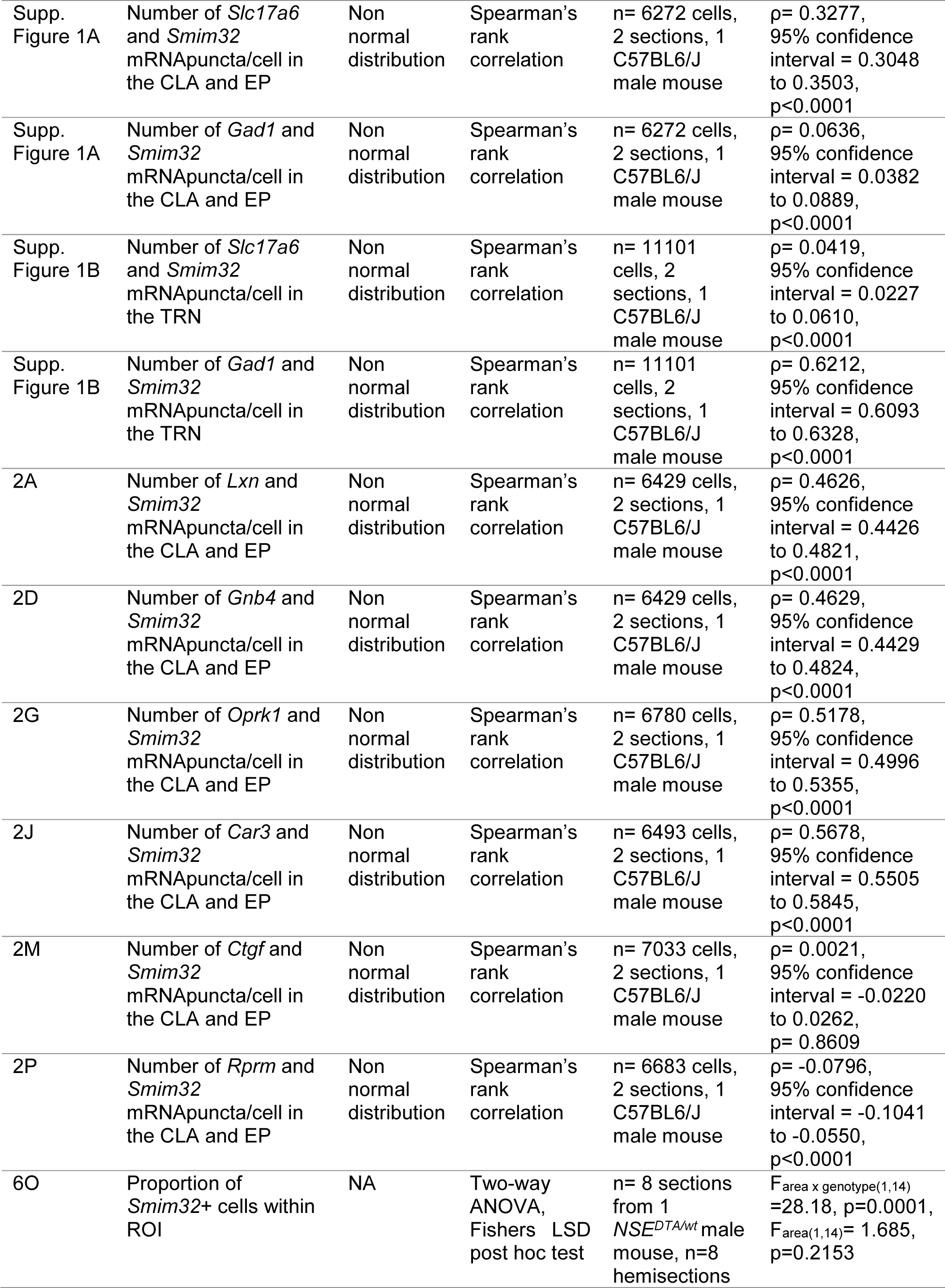
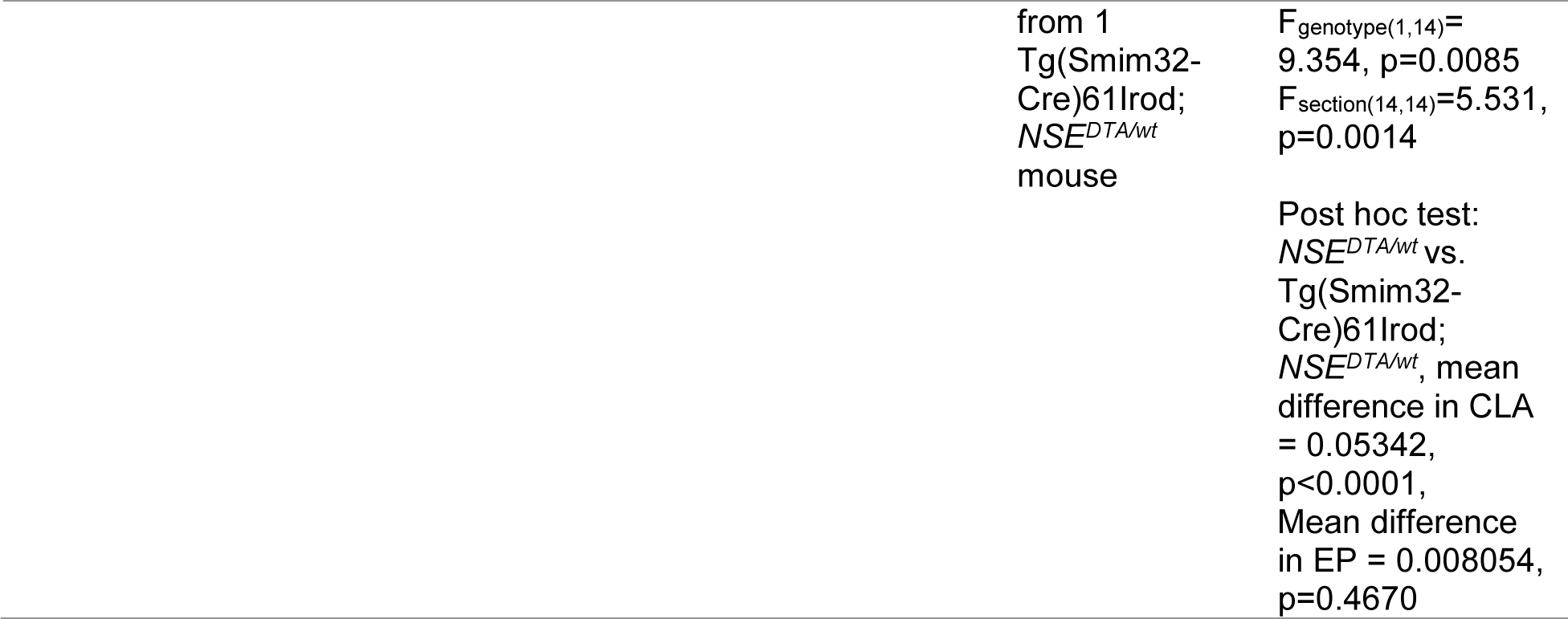
Statistical data.

### RNAscope image analysis

Images were analyzed with QuPath, version 0.4.3. First, cell segmentation based on DAPI staining was performed using the *Cell detection* function. Nuclei with a total area below 30 μm^2^ were excluded from the analysis. To include transcripts located in the cytoplasm, the area of the cells was expanded by 2 μm around the nucleus. Next, puncta corresponding to individual mRNA molecules were automatically detected and quantified using the *Subcellular detection* function. For each probe/fluorophore, a detection threshold is set, and all puncta above this threshold are counted. The same threshold is applied to all images labeled with a specific probe. When multiple puncta are clustered together, an estimation of the number of individual puncta within the cluster is performed automatically based on the intensity and size of the cluster.

#### Co-expression analysis

Cells were considered positive for a gene expression when a minimum of 20 mRNA puncta of the corresponding probe were attributed to this cell. This threshold was optimized in order to exclude false positives. The threshold was adapted to a minimum of 5 puncta/cell when the overall gene expression was very low in the cell population (the case for *Car3* and *Slc17a6*).

### Cre/FlpO reporter imaging and analysis

Mice heterozygous for *Smim32-Cre* or hemizygous for Tg(Smim32-Cre)61Irod or Tg(Smim32-Cre)62Irod were crossed with a reporter mouse line homozygous for *Rosa26-stop(LoxP)-tdTomato,* to generate heterozygous reporter mice. Expression of tdTomato in Smim32-Cre male and female mice were analyzed at age E16, E18, P0, P2, P4, P6, P8, P10, P12, P14, P16, P18, P20, P70 and P200. Tg(Smim32-Cre)61Irod and Tg(Smim32-Cre)62Irod male mice were analyzed at age P0, P2, P4, P6, P8, P10, P12, P14, P16, P18, P20 and P70. A male Smim32-FlpO mouse was analyzed at P20. One mouse per time point, sex and genotype were analyzed.

#### Embryos (E16-E18) and Newborns (P0-P8)

To collect embryos, pregnant mothers received in intra peritoneal (i.p.) injection of the opioid analgesic buprenorphine at 0.1 mg/kg (Temgesic, Eumedica Pharmaceuticals AG). After 30 minutes, they received a lethal i.p. injection of pentobarbital at 150 mg/kg (Esconarcon, Streuli Pharma SA). Mice were perfused transcardially with heparin and then with formalin 10%. After perfusion, embryos were collected and fixed overnight in formalin 10% at 4°C. For newborns up to P6, pups were decapitated and heads were fixed in formalin 10% overnight at 4°C. P8 mice were euthanized as described for pregnant mothers, before perfusion with heparin and formalin 10%, followed by immersion in formalin 10 overnight at 4°C. Heads were immersed in PBS 15% sucrose for 8 hours and PBS 30% sucrose overnight at 4°C, embedded in OCT and frozen in an isopentane bath in liquid nitrogen. Heads were cut on a cryostat (Leica DM 1850) in 20 μm sections and stored at -80°C until analysis. Before analysis, slides were stained with DAPI (1 μg/ml in PBS) and mounted with DABCO (25 mg/ml) in a mix of 90% glycerol 10% water.

#### Young and adult mice (P10-P200)

Mice were euthanized as described for pregnant mothers. After perfusion, brains were collected and fixed overnight in formalin 10% at 4°C. Brains were embedded in 3% low melt agarose (TopVision Low Melting Point Agarose, Thermo Fisher Scientific®) and cut in 50 μm sections using a vibratome (Microm HM 650 V). Sections stained with DAPI, placed on Superfrost Plus slides, and mounted with DABCO.

#### Imaging

Slides were imaged on an epifluorescence Leica DM5500B microscope with a HC PL FLUOTAR 10x/0.30 DRY objective and a Leica Leica-K5 camera. Image acquisition and merging of tile scanc was done using the LAS X software (version 3.7.4.23463). Tile scans of entire coronal sections were acquired for UV (470 nm), Cy3 (605 nm) and GFP (527 nm) channels.

#### Image analysis

In embryos and young mice up to P8, the expression of tdTomato was assessed visually and manually. Once multiple cells could be clearly identified within a structure, on several consecutive sections and bilaterally, expression onset was noted. After P8, relative expression levels of tdTomato were quantified by counting the proportion of DAPI-positive cells expressing tdTomato within specific brain areas. Image analysis was done using QuPath (version 0.4.3). Cells were segmented passed on DAPI staining. Next, tdTomato-expressing cells were identified using the Subcellular detection tool, set to detect both red (tdTomato) and green (autofluorescence) signals. All cells with autofluorescence signals were excluded from the analysis.

### IHC on free floating sections

Mice were anesthetized by an i.p. injection of pentobarbital (200 mg/kg) and perfused transcardially with heparin and then with formalin 10%. Brains were kept overnight in the same fixative at 4°C and then embedded in agarose low melt (Thermo Fisher Scientific®) to be cut in 50 μm sections using a vibratome. Slices were post-fixed 10 minutes in formalin 10% at room temperature and washed in 1x PBS. Sections were pre-incubated in incubation solution (1x PBS, Triton X-100 0.5%, FCS 10%) for 1 hour at room temperature with agitation. Primary and secondary antibodies were diluted in incubation solution. Sections were incubated with primary antibody overnight at 4°C with agitation. Sections were washed 3×15 in washing solution (1x PBS Triton X-100 0.5%) before incubating for 3 hours at room temperature with a secondary antibody. Sections were washed 3×15 in washing solution, and nuclei were stained with DAPI (1 μg/ml in PBS) for 5 minutes at room temperature. Slides were mounted with DABCO (25 mg/ml) in a mix of 90% glycerol and 10% water. The double IHC was performed similarly but mixing both primary and secondary antibodies. When only endogenous fluorescence was observed, mice were prepared as for IHC on free-floating sections, but without any incubations except DAPI.

### IHC antibodies

**IHC imaging** (Figure 6)

#### Fluorescence microscopy

Tile scans of coronal sections were made using a Leica DM5500 microscope. Stitching was performed with the build-in Tilescan module of the LAS X software. For expression screening experiments, specific tdT fluorescence was controlled by verifying the absence of autofluorescence in the green channel.

#### Confocal microscopy

Confocal images were acquired using a Leica SP8 confocal microscope and deconvolved with the build-in Hyvolution module.

**Viral injections** (Figure 6)

#### Injection

A 4-month-old Tg(Smim32-Cre)61Irod female mouse was anesthetized with isoflurane (3–5% induction, 1–2% maintenance), and the skin overlaying the skull was removed under local anesthesia. Mice were then head-fixed with ear bars and a nose clamp on a stereotaxic apparatus (Stoelting). Eyes were protected from drying with artificial tears. The body temperature was monitored with a rectal probe and was maintained at ∼37°C using a heating pad (FHC) during surgery. Bilateral craniotomies were performed above the CLA using an air-pressurized driller. Glass pipettes filled with solutions containing AAV2-Ef1a-DIO-hChR2-(E123/T159C)-EYFP were inserted in one hemisphere at a time. Antero-posterior (AP): 1.1 mm, medio-lateral (ML): ± 2.85 mm relative to bregma and dorso-ventral (DV): 2.4 mm relative to brain surface. A volume of ∼100 nL was slowly injected. The pipette was left in place for a period of 3-5 min before its removal and the suturing of the skin. All infections were bilateral unless specified.

#### Analysis of viral expression

For evaluation of viral expression in the CLA, the mouse was anesthetized ∼3 weeks later by an intraperitoneal (i.p.) injection of pentobarbital (200 mg/kg) and perfused transcardially with 20 ml of phosphate-buffered saline (PBS, 0.1 M, pH 7.3) followed by 50 ml of 4% paraformaldehyde (PFA) in 0.1 M PBS at 4 °C. The brain was removed and left overnight in 4% PFA. After embedding the brains in 4% agarose, 40 μm coronal slices were cut with a vibratome and collected in PBS (0.1 M). For immunostaining, slices were rinsed in PBS 0.1% TX-100, 0.7% H2O2. Slices were then incubated with PBS 1% BSA for 1 hour at 21-25 °C, and then with the primary antibody rabbit anti-YFP (1:1000, Invitrogen, ref. A6455), overnight at 4 °C. The day after, slices were rinsed and incubated with biotinylated goat anti-rabbit IgG (1:200, Vector, ref. BA-1000) for 1 hour at room temperature. Slices were then processed using an avidin– biotin– peroxidase complex (ABC kit, Vector Laboratories) and reacted with the chromogen 3,3’-diaminobenzidine (DAB, Sigma-Aldrich). Slices were mounted with Aquatex mounting medium.

### Statistics

## ACKNOWLEDGMENTS

We thank Véronique Pauli, Chenda Kan, Enes Karavdic and Francisco Resende for expert technical assistance. We thank all members of I.R and A.C laboratories helpful discussions and for providing examples of usage of our transgenic mouse lines. We thank the Transgenesis Core Facility of the University of Geneva and the Center for Transgenic Models of the University of Basel for assistance in generating the transgenic mouse lines.

## COMPETING INTERESTS

The authors declare that they have no competing interests.

## FUNDING

This research was supported by the University of Geneva and the European Research Council (contract ERC-SyG-856439-CLAUSTROFUNCT to A.C. and I.R.), the Swiss National Science Foundation (grant 310030_215572 to and 310030_219531 to A.C. and I.R., respectively), the Fondation Privée des HUG (A.C. and I.R.), and the Novartis foundation for medical research (A.C. and I.R.).

## AUTHOR CONTRIBUTIONS

Ivan Rodriguez and Alan Carleton, conceptualization, funding acquisition, supervision, writing – original draft, review & editing; Joël Tuberosa and Madlaina Boillat, investigation, project administration, formal analysis, visualization, writing – original draft; Julien Dal Col, methodology; Leonardo Marconi, investigation, formal analysis; Julien Codourey, Loris Mannino, Elena Georgiou, Marc Menoud, investigation.

## SUPPLEMENTARY FIGURES

**Supplementary Figure 1.**
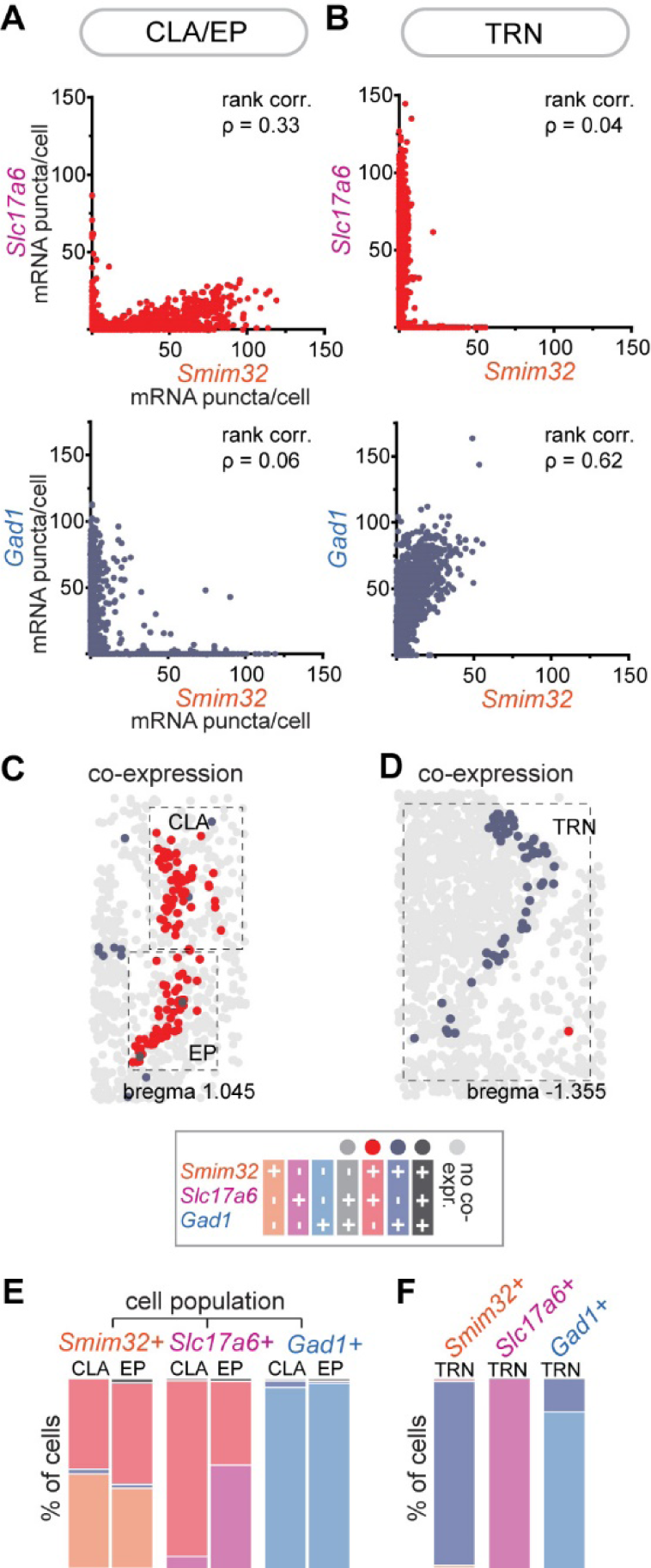
*Smim32* is expressed in excitatory neurons in the CLA and in inhibitory neurons in the TRN. (**A,B**) Correlation between expression levels of *Smim32* and *Slc17a6, and Smim32* and *Gad1,* in cells from the (**A**) CLA/EP and from the (**B**) TRN. Each dot corresponds to one cell. All cells from both hemispheres, within the regions of interest (ROI, dotted line in (**C,D**)), were included in the analysis. ρ value indicates Spearman’s rank correlation. CLA/EP, n = 6272 cells; TRN, n = 11101 cells. (**C,D**) Illustrative map of cells co-expressing *Smim32* with *Slc17a6* (red) or *Smim32* with *Pvalb* (purple grey) in the CLA/ EP and in the TRN. A cell was considered positive for a marker when a minimum of 20 mRNA puncta were attributed to this cell. The dotted lines highlight the ROIs used for quantification analyses shown in (**E,F**). (**E,F**) Bar plots showing the percentages of cells positive for one or several markers. The analysis was done separately for each cell population.

**Supplementary Figure 2.**
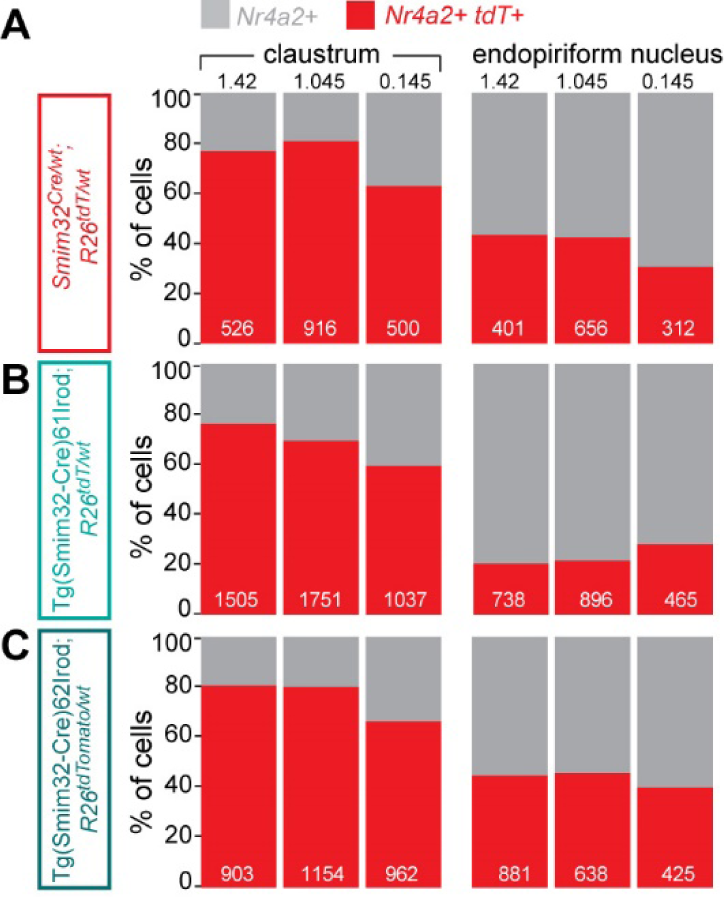
Proportion of CLA neuronal population expressing Cre driven reporter. (**A**) Quantification of *tdT* expressing cells in *Nr4a2*-positive neuronal population in *Smim32^Cre/wt^; R26^tdT/wt^* mice. Bar plots show the percentage of *Nr4a2*-positive cells from the CLA and EP that co-express *tdT*. Numbers in the bar plots indicate the number of cells analyzed in the corresponding region and anteroposterior position. n= 1 *Smim32^Cre/wt^; R26^tdT/wt^* mouse. (**B-C**) Same as above, but for Tg(Smim32-Cre)61Irod and Tg(Smim32-Cre)62Irod mice. n= 2 Tg(Smim32-Cre)61Irod mice, n=1 Tg(Smim32-Cre)62Irod mouse.

**Supplementary Figure 3.**
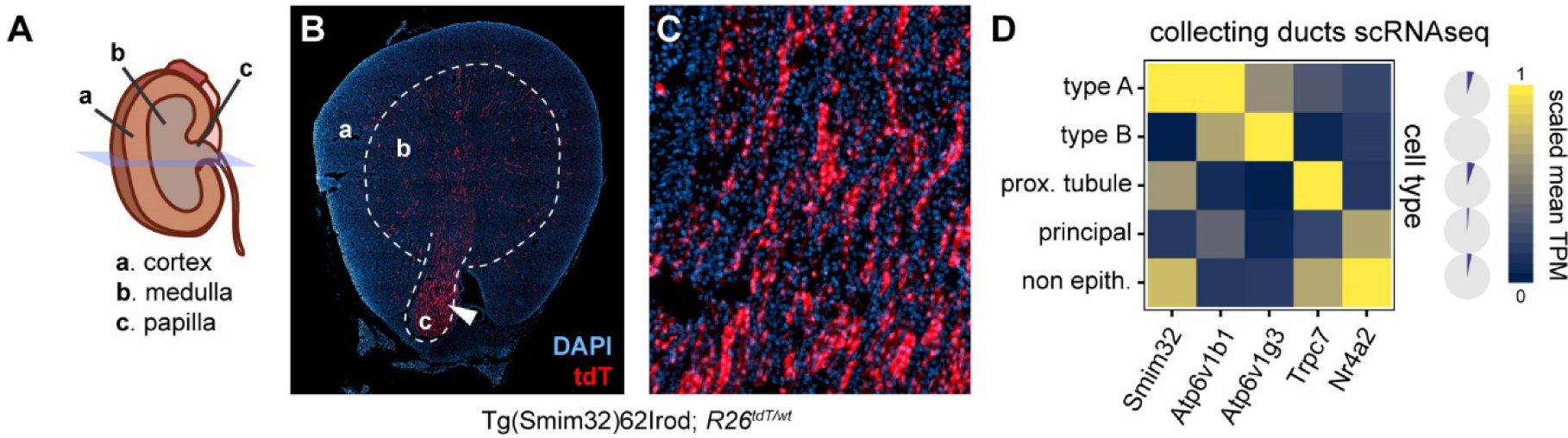
*Smim32* expression in the kidney. (**A**) Schematic of a mouse kidney with the cutting plane of the section shown in (**B**) and (**C**). (**B**) Reporter expression in the kidney of a Tg62(Smim32-Cre) mouse. (**C**) Zoom on the papilla, where arrays of reporter-expressing cells can be observed, likely forming the walls of distal tubules. (**D**) Re-analysis of mouse collecting duct scRNAseq data from Chen et al^40^. Statistics were computed for cell types (rows of the matrix) and genes of interest (columns of the matrix). For each gene, mean expression values (mean TPM) were divided by the maximum observed mean expression across all samples to produce the scaled mean TPM value (max mean TPMs: *Smim32*=0.216, *Atp6v1b1*=353, *Atp6v1g3*=3088, *Trpc7*=2.06, *Nr4a2*=8.35). On the right, the dark areas of the pie charts represent the proportion of cells expressing more than 1 TPM of *Smim32* for each cell type.

**Supplementary Figure 4.**
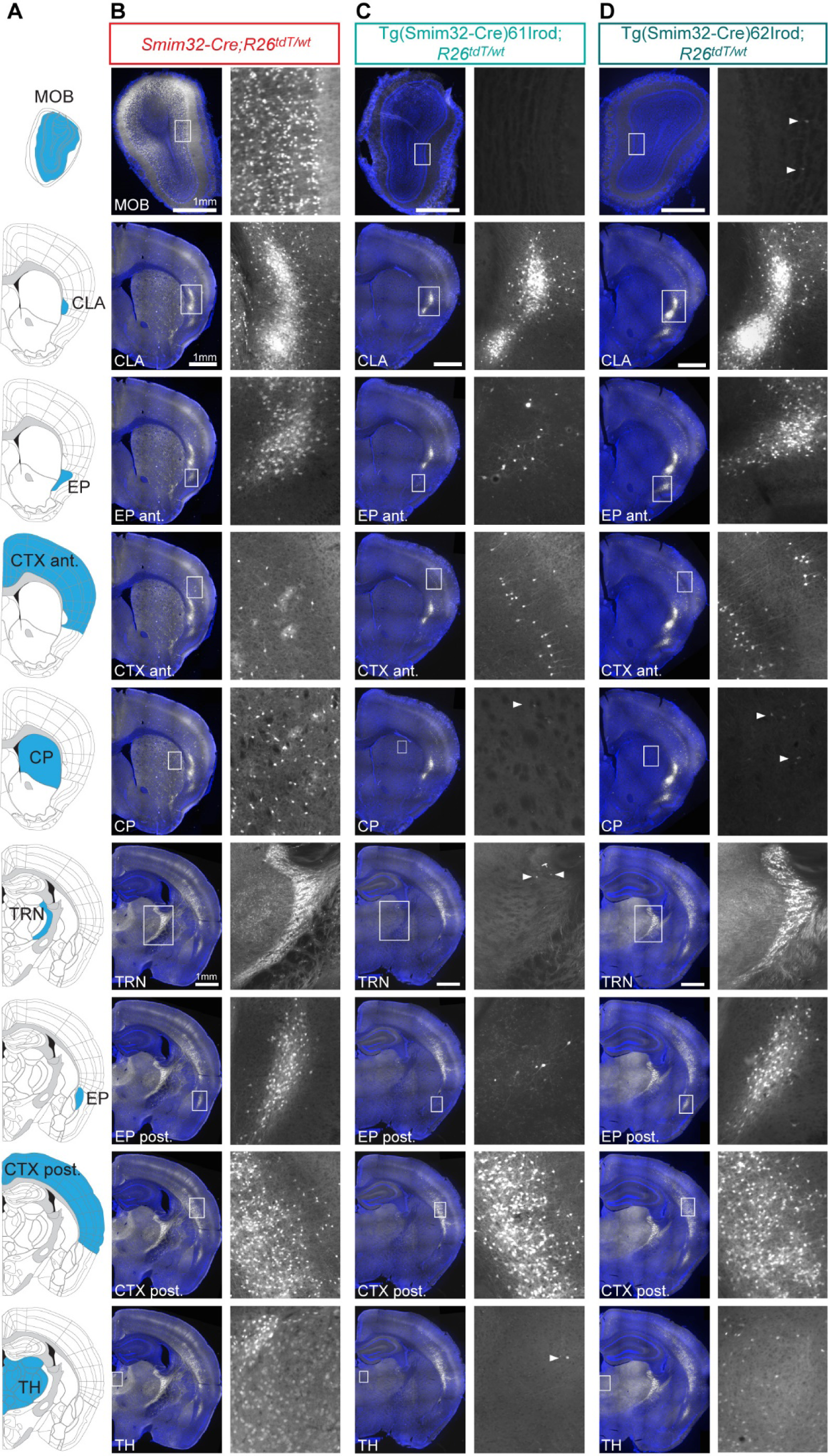
Representative images of Cre-driven tdT expression in *Smim32* transgenic mice. (**A**) Schematics of the regions of interest within which tdT expression was quantified. (**B-D**) Representative epifluorescence images of tdT expression in *Smim32-Cre,* Tg(Smim32-Cre)61Irod *and* Tg(Smim32-Cre)62Irod mice carrying the Cre-dependent *R26-tdT* allele. Left panels, overview of the section stained with DAPI (blue) and endogenous tdT fluorescence (white). Scale bar: 1 mm. The white square highlights the region magnified in the right panels. White arrowheads point towards tdT expressing cells in regions where expression is very sparse.

## Notes

### Competing Interest Statement

The authors have declared no competing interest.

## REFERENCES

1. Wang, Q. et al. Organization of the connections between claustrum and cortex in the mouse. J. Comp. Neurol. 525, 1317–1346 (2017).

2. Zingg, B. et al. Neural Networks of the Mouse Neocortex. Cell 156, 1096–1111 (2014).

3. Torgerson, C. M., Irimia, A., Goh, S. Y. M. & Van Horn, J. D. The DTI connectivity of the human claustrum: Claustrum Connectivity. Hum. Brain Mapp. 36, 827–838 (2015).

4. Marriott, B. A., Do, A. D., Zahacy, R. & Jackson, J. Topographic gradients define the projection patterns of the claustrum core and shell in mice. J. Comp. Neurol. (2020) doi:10.1002/cne.25043.

5. Chia, Z., Augustine, G. J. & Silberberg, G. Synaptic Connectivity between the Cortex and Claustrum Is Organized into Functional Modules. Curr. Biol. 30, 2777–2790.e4 (2020).

6. Shelton, A. M. et al. Single neurons and networks in the claustrum integrate input from widespread cortical sources. Preprint at 10.1101/2022.05.06.490864 (2022).

7. Ham, G. X. & Augustine, G. J. Topologically Organized Networks in the Claustrum Reflect Functional Modularization. Front. Neuroanat. 16, 901807 (2022).

8. Wang, Q. et al. Regional and cell-type-specific afferent and efferent projections of the mouse claustrum. Cell Rep. 42, 112118 (2023).

9. Crick, F. C. & Koch, C. What is the function of the claustrum? Philos. Trans. R. Soc. Lond. B. Biol. Sci. 360, 1271–1279 (2005).

10. Koubeissi, M. Z., Bartolomei, F., Beltagy, A. & Picard, F. Electrical stimulation of a small brain area reversibly disrupts consciousness. Epilepsy Behav. EB 37, 32–35 (2014).

11. Remedios, R., Logothetis, N. K. & Kayser, C. Unimodal Responses Prevail within the Multisensory Claustrum. J. Neurosci. 30, 12902–12907 (2010).

12. Remedios, R., Logothetis, N. K. & Kayser, C. A role of the claustrum in auditory scene analysis by reflecting sensory change. Front. Syst. Neurosci. 8, (2014).

13. Smythies, J., Edelstein, L. & Ramachandran, V. Hypotheses relating to the function of the claustrum. Front. Integr. Neurosci. 6, (2012).

14. Grasby, K. & Talk, A. The anterior claustrum and spatial reversal learning in rats. Brain Res. 1499, 43–52 (2013).

15. Fodoulian, L., et al. The Claustrum-Medial Prefrontal Cortex Network Controls Attentional Set-Shifting. http://biorxiv.org/lookup/doi/10.1101/2020.10.14.339259 (2020) doi:10.1101/2020.10.14.339259.

16. White, M. G. et al. The Mouse Claustrum Is Required for Optimal Behavioral Performance Under High Cognitive Demand. Biol. Psychiatry 88, 719–726 (2020).

17. Atlan, G. et al. The Claustrum Supports Resilience to Distraction. Curr. Biol. 28, 2752–2762.e7 (2018).

18. Narikiyo, K. et al. The claustrum coordinates cortical slow-wave activity. Nat. Neurosci. 23, 741–753 (2020).

19. Norimoto, H. et al. A claustrum in reptiles and its role in slow-wave sleep. Nature 578, 413–418 (2020).

20. Renouard, L. et al. The supramammillary nucleus and the claustrum activate the cortex during REM sleep. Sci. Adv. 1, e1400177 (2015).

21. Kitanishi, T. & Matsuo, N. Organization of the Claustrum-to-Entorhinal Cortical Connection in Mice. J. Neurosci. 37, 269–280 (2017).

22. Dixsaut, L. & Gräff, J. Brain-wide screen of prelimbic cortex inputs reveals a functional shift during early fear memory consolidation. eLife 11, e78542 (2022).

23. Mutel, S., Renfer, Jr., Rodriguez, I., Carleton, A. & Salazar, Rf. The Claustrum Drives Large-Scale Interactions of Cortical Circuits Relevant to Long-Term Memory. http://biorxiv.org/lookup/doi/10.1101/2023.03.02.530783 (2023) doi:10.1101/2023.03.02.530783.

24. White, M. G. et al. Anterior Cingulate Cortex Input to the Claustrum Is Required for Top-Down Action Control. Cell Rep. 22, 84–95 (2018).

25. Liu, J. et al. The Claustrum-Prefrontal Cortex Pathway Regulates Impulsive-Like Behavior. J. Neurosci. 39, 10071–10080 (2019).

26. Atlan, G., et al. Claustral Projections to Anterior Cingulate Cortex Modulate Engagement with the External World. http://biorxiv.org/lookup/doi/10.1101/2021.06.17.448649 (2021) doi:10.1101/2021.06.17.448649.

27. Chevée, M., Finkel, E. A., Kim, S.-J., O’Connor, D. H. & Brown, S. P. Neural activity in the mouse claustrum in a cross-modal sensory selection task. Neuron 110, 486–501.e7 (2022).

28. Jackson, J., Karnani, M. M., Zemelman, B. V., Burdakov, D. & Lee, A. K. Inhibitory Control of Prefrontal Cortex by the Claustrum. Neuron 99, 1029–1039.e4 (2018).

29. Xu, Q.-Y. et al. Identification of a Glutamatergic Claustrum-Anterior Cingulate Cortex Circuit for Visceral Pain Processing. J. Neurosci. 42, 8154–8168 (2022).

30. Koga, K., Kobayashi, K., Tsuda, M., Pickering, A. E. & Furue, H. Anterior cingulate cross-hemispheric inhibition via the claustrum resolves painful sensory conflict. *Commun*. Biol. 7, 330 (2024).

31. Ntamati, N. R., Acuña, M. A. & Nevian, T. Pain-induced adaptations in the claustro-cingulate pathway. Cell Rep. 42, 112506 (2023).

32. Terem, A. et al. Claustral Neurons Projecting to Frontal Cortex Mediate Contextual Association of Reward. Curr. Biol. 30, 3522–3532.e6 (2020).

33. Niu, M. et al. Claustrum mediates bidirectional and reversible control of stress-induced anxiety responses. Sci. Adv. 8, eabi6375 (2022).

34. Yao, Z. et al. A taxonomy of transcriptomic cell types across the isocortex and hippocampal formation. Cell 184, 3222–3241.e26 (2021).

35. Peng, H. et al. Morphological diversity of single neurons in molecularly defined cell types. Nature 598, 174–181 (2021).

36. Daigle, T. L. et al. A Suite of Transgenic Driver and Reporter Mouse Lines with Enhanced Brain-Cell-Type Targeting and Functionality. Cell 174, 465–480.e22 (2018).

37. Ibrahim, C., Le Foll, B. & French, L. Transcriptomic Characterization of the Human Insular Cortex and Claustrum. Front. Neuroanat. 13, 94 (2019).

38. Zeisel, A. et al. Molecular Architecture of the Mouse Nervous System. Cell 174, 999–1014.e22 (2018).

39. Madisen, L. et al. A robust and high-throughput Cre reporting and characterization system for the whole mouse brain. Nat. Neurosci. 13, 133–140 (2010).

40. Chen, L. et al. Transcriptomes of major renal collecting duct cell types in mouse identified by single-cell RNA-seq. Proc. Natl. Acad. Sci. 114, (2017).

41. Osoegawa, K. et al. Bacterial artificial chromosome libraries for mouse sequencing and functional analysis. Genome Res. 10, 116–128 (2000).

42. Sando, R., 3rd et al. Inducible control of gene expression with destabilized Cre. Nat Methods 10, 1085–8 (2013).

43. Yao, X. et al. Tild-CRISPR Allows for Efficient and Precise Gene Knockin in Mouse and Human Cells. Dev. Cell 45, 526–536.e5 (2018).

44. Butler, A., Hoffman, P., Smibert, P., Papalexi, E. & Satija, R. Integrating single-cell transcriptomic data across different conditions, technologies, and species. Nat. Biotechnol. 36, 411–420 (2018).

45. Dobin, A. et al. STAR: ultrafast universal RNA-seq aligner. Bioinformatics 29, 15–21 (2013).

46. Liao, Y., Smyth, G. K. & Shi, W. featureCounts: an efficient general purpose program for assigning sequence reads to genomic features. Bioinforma. Oxf. Engl. 30, 923–930 (2014).

